# Charting the liver and lung metastatic niche in breast cancer

**DOI:** 10.1101/2024.10.04.616491

**Authors:** Magdalena K. Sznurkowska, Francesc Castro-Giner, Ilona Krol, Yongzhan Zhang, Lauren L. Ozimski, Irene D’Anna, Alexander Ring, Fabienne Dominique Schwab, Yu Wei Zhang, Jianwen Zhou, Massimo Saini, Karin Strittmatter, Selina Budinjas, Marko Vujanovic, Zacharias Kontarakis, Giada Pontecorvi, Francesca Albrecht, Kirsten D. Mertz, Gaël Auray, Claudio Giachino, Verdon Taylor, Ana Gvozdenovic, Werner J. Kovacs, Ilaria Malanchi, Heike Frauchiger-Heuer, Andreas Wicki, Marcus Vetter, Matteo Ligorio, Nicola Aceto

## Abstract

Breast cancer progression to visceral organs such as lung and liver is regarded as a dreadful event, unequivocally associated with a poor prognosis. Yet, these vital sites are characterized by highly diverse cellular microenvironments and physiological functions, suggesting that they may influence cancer cells behavior in divergent ways. Unexpectedly, we find that while the liver microenvironment fosters metastasis-promoting properties and boosts secondary spread, the lungs impose a roadblock to the same processes. Using patient data and tissues from rapid autopsy, as well as mouse models with barcode-mediated metastasis tracing, niche labeling technology and single cell analysis of both tumor cells and their direct microenvironment, we dissect cellular and molecular microenvironmental factors that impose this differential behavior. Among these, we identify BMP2-producing endothelial cells as critical players within the liver metastatic niche, capable to enhance metastasis-to-metastasis dissemination. Targeting BMP2 receptor on breast cancer cells suppresses their metastasis-forming ability. Altogether, we reveal a contrast in the site-specific behavior of lung and liver metastases in breast cancer, highlighting microenvironmental factors that contribute to this diversity, as well as organ-specific opportunities for intervention.

## Main

The spread of cancer is a major cause of cancer-related death (*1*). While the newest emerging treatments are typically tested in patients with a metastatic disease, most of the underlying mechanistic insights that led to their unfolding derive from explorations of primary tumor biology (*2–4*). Consequently, little is known about how cancer cells are shaped at different anatomical sites upon dissemination. A better understanding of cancer cell biology as a function of their location is key to expose site-specific vulnerabilities and tailor new treatment interventions to a metastatic disease.

Breast cancer is the most common malignancy in women (*5*), typically progressing to bone, liver, lungs and brain (*6*). The triple-negative breast cancer (TNBC) subtype, defined by the lack of expression of HER2, estrogen and progesterone receptors (ER, PR), amounts to 10-15% of total breast cancers (*7*, *8*) and is characterized by an exceptionally aggressive behavior, limited treatment opportunities and consequently, poor outcome compared to other subtypes (*7*, *8*). Intriguingly, TNBC spreads primarily to visceral organs, most commonly lungs and liver (*9*), in contrast to hormone receptor positive breast cancers that display a more pronounced tropism for the bone (*10–13*). These metastatic patterns appear to affect disease outcome. For instance, the presence of bone metastases is associated with a more indolent disease and long-term outcomes, compared to other sites (*14*, *15*). While reports on disease aggressiveness in the context of visceral metastases in the lungs and liver vary (*11*, *12*, *15*), these two organs display substantially different anatomical and physiological features, suggesting that they may impact on cancer cell biology in a different way.

Based on this, we hypothesized that lungs and liver shape breast cancer metastases differently, that the components of the metastatic niche within these organs vary, and that metastasis location could have direct impact on progression-related features such as likelihood of secondary seeding. We sought to uncover unique, site-specific biological changes that expose cancer cells to “geographical” vulnerabilities, suggesting novel therapies that are designed not only based on general cancer-associated features, but also on disease location.

### Site-specific impact on survival

We first sought to establish whether differential metastatic patterns could affect patient survival in breast cancer patients. To this end, we examined the clinical history of 318 patients diagnosed with metastatic breast cancer between 1972 and 2019 at the University Hospital Basel and University Hospital Zurich, Switzerland. Compared to most public cancer patient databases that collect recurrence data indirectly (*16*), our cohort contained direct diagnostic (metastatic) follow up information, allowing us to precisely define the metastatic progression sequence of each patient (**Supplementary Tables 1,2**). With the aim to assess whether metastasis to liver or lung could have a differential impact on survival, we focused on rare cases with mono-metastasis (*n* = 22), whereby a single metastasis in lung (*n* = 12) or liver (*n* = 10) was present at the initial metastatic diagnosis (**Fig. 1A**). Of note, in this cohort, while concurrent liver and lung metastasis were generally prevalent as expected (64.2% and 55.3%, respectively), the occurrence of a single, visceral metastasis as first metastatic diagnosis was noticeably rarer (3.8% for lung, 3.1% for liver). We found that patients with liver metastasis were characterized not only by a significantly shorter overall survival (Cox Proportional Hazard (PH) *P* value = 0.022; hazard ratio (HR) = 3.3, 95% confidence interval (CI), 1.19 to 9.10), consistently with a previous report (*11*), but also a shorter time to next metastatic event or death (Cox PH *P* value = 0.006; HR = 9.4, 95%CI, 1.89 to 46.89) compared to patients with lung metastases (**Fig. 1B**). Of note, and in line with previous observations (*14*, *15*), metastases to the bone were associated with a better prognosis, comparable to lung metastases (bone vs. liver: Cox PH *P* value = 0.00048 and HR = 0.23, 95%CI, 0.15 to 0.53; bone vs. lung: Cox PH *P* value = 0.23 and HR = 1.49, 95%CI, 0.78 to 2.86; **Extended Data** Fig. 1A, B). We then tested in an independent, external dataset from Surveillance, Epidemiology, and End Results Program (SEER) (*17*), whether the presence of liver metastases was also associated with a poorer prognosis compared to lung metastasis in TNBC. Also in this case, we found that the time to next metastasis or death was longer for patients with initial lung metastasis diagnosis (Cox PH *P* value = 0.0095 and HR = 0.82, 95%CI, 0.71 to 0.95; **Extended Data** Fig. 1C). Possibly underlying a general metastasis feature, a significant association was also observed in other cancer types, with the time until next metastasis being shorter for colorectal and pancreatic ductal adenocarcinoma (PDAC) cancer patients with an initial liver metastasis diagnosis (colorectal: Cox PH *P* value = 6.53e-09 and HR = 0.85, 95%CI, 0.81 to 0.90, PDAC: Cox PH *P* value < 2.22e-16 and HR = 0.84, 95%CI, 0.81 to 0.87; **Extended Data** Fig. 1D,E). Altogether, these results highlight an intriguing association between survival probability and organ-specific metastasis within visceral sites, suggesting an accelerated disease progression in patients with a first visceral metastasis within the liver.

**Fig. 1.**
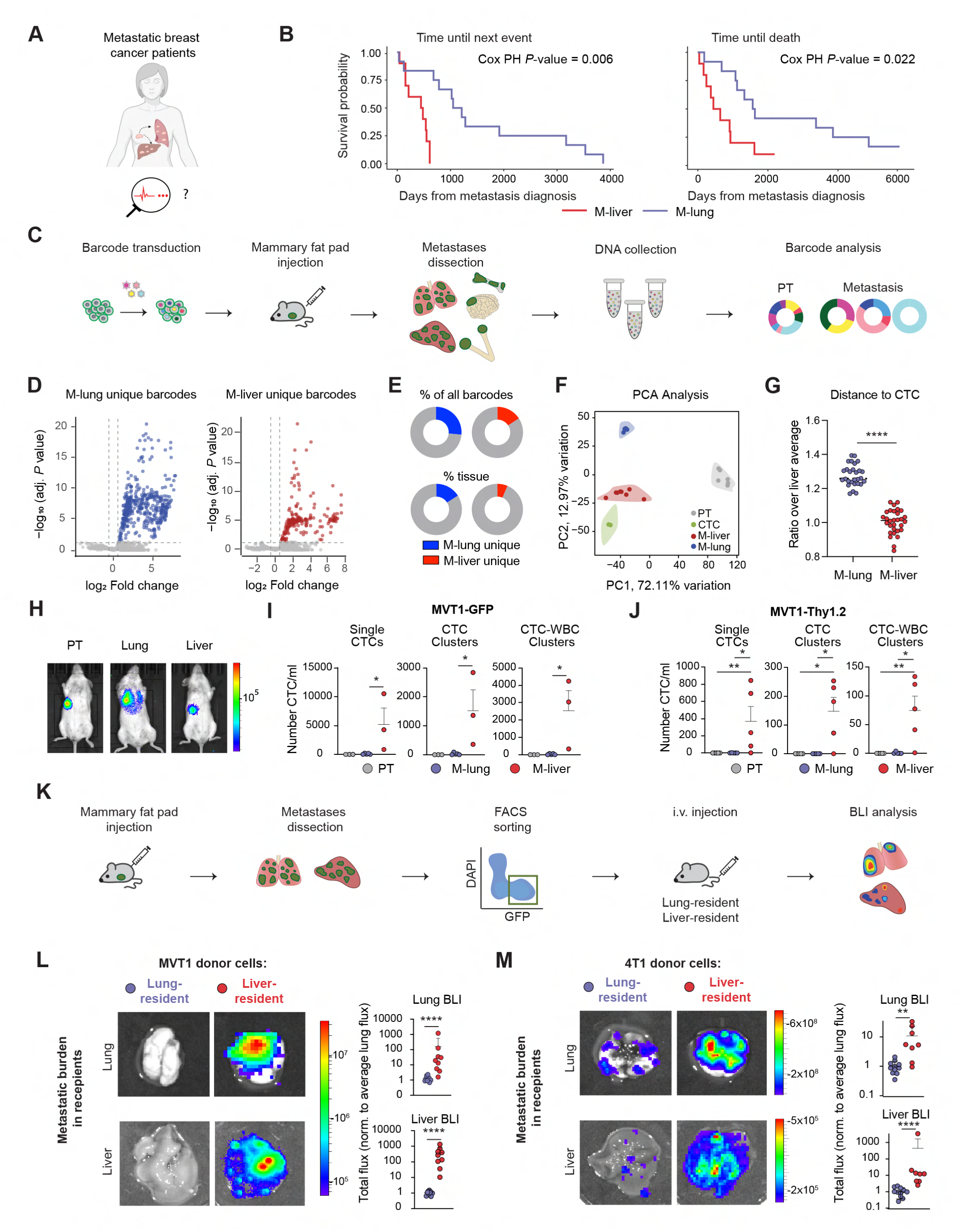
Metastatic ability of liver-resident *versus* lung-resident breast cancer cells. **A**. Schematic showcasing the concept behind patient data analysis. **B.** Kaplan-Meier plots depicting time to next event (*left*) and overall survival (*right*) of breast cancer patients with the first mono-metastatic event detected in lungs (*n* = 12) or liver (*n* = 10). **C.** Schematic representing the experimental design for barcode labelling and metastasis clonality assessment in NSG mice carrying MVT1-GFP-luciferase tumors. **D.** Volcano plots illustrating unique barcodes for lung (*left*, M-lung) and liver (*right*, M-liver) metastases of a representative mouse. **E.** Pie charts showing the percent of unique barcodes detected in lungs or liver metastases (*top*), as well as the unique barcode contribution to metastasis in lungs and liver (*bottom*, M-lung and M-liver). **F.** Principal component analysis (PCA) plot of 500 top variable barcodes represented in the primary tumor (PT), circulating tumor cells (CTCs), lung and liver metastases of a representative mouse. **G.** Dot plot showing Euclidean distances measuring the dissimilarity in barcode expression between lung or liver metastases and CTCs (*n* = 3 mice). ****, *P* < 0.0001 by Mann-Whitney test. **H.** Representative bioluminescence images of mice following injections into mammary fat pad (*left*), intravenous injection into lungs (*center*) and splenic injection into liver (*right*). **I-J**. Enumeration of single CTCs, CTC clusters, and CTC-white blood cell (WBC) clusters for the MVT1-GFP in NSG model (**I,** *n* = 3 mice for primary tumor and liver, and *n* = 5 mice for lung) and the MVT1-Thy1.2 in FVB model (**J,** *n =* 5 mice per group) two weeks after mammary fat pad injection (PT), i.v. injection (M-lung) and intrasplenic injection (M-liver). **, *P* < 0.01, *, *P* < 0.05 by Mann-Whitney test. **K.** Illustration of the experimental design for metastatic potential assessment of lung- and liver-resident breast cancer cells. **L-M.** Representative images (*left*) and quantification (*right*) of bioluminescent signals in indicated organs of animals injected with lung- or liver-resident MVT1-GFP-luciferase (**L**, *n* = 9 mice per group) or 4T1-GFP-luciferase cells (**M,** *n* = 11 mice per lung-derived, *n* = 8 mice per liver-derived groups). **, *P* < 0.01, ****, *P* < 0.0001 by Mann-Whitney test. i.v., intravenous; BLI, bioluminescence imaging. Panel **A** was created with BioRender.com.

### Clonal tracking of visceral metastases *in vivo*

We next sought to verify whether the correlation observed with the specific visceral site and disease aggressiveness could be attributed to the functional properties of the cells residing in the respective organs using a TNBC mouse model. To this end, we employed cellular barcoding, whereby MVT1 breast cancer cells were uniquely labelled with a genetic barcode library composed of five million barcodes. After barcoding, 100’000 MVT1 cells were injected orthotopically into the mammary fat pad of NOD.Cg-*Prkdc*^scid^-*Il2rg*t^m1Wjl^/SzJ (NSG) female mice, mimicking physiological selection of the fittest clones as well as adaptation at target metastatic sites upon spontaneous dissemination. Of note, we purposely did not use immunocompetent syngeneic models for these experiments, to avoid well-known immune system-mediated rejection of non-self immunogenic components such as Cas9, barcode-associated fluorescence markers and genes conferring resistance to substances with eukaryotic toxicity (*18*). Upon spontaneous metastasis to visceral sites, we performed blood draw for circulating tumor cell (CTC) isolation as well as dissection of both primary and metastatic sites, followed by targeted barcode sequencing to determine clonal composition (**Fig. 1C**). Using a cutoff of five counts per million, we identified on average 11’091 barcodes in the primary tumor, 1’305 barcodes in CTCs, 918 and 1’094 barcodes in liver and lung metastases, respectively (**Extended Data** Fig. 2A). We reasoned that the frequency and abundance of barcodes unique to a particular metastatic site would reflect a limited ability to achieve secondary seeding. When focusing on unique barcodes in lungs and liver with respect to all the other metastatic sites (**Extended Data** Fig. 2B-D), we observed that they were substantially more frequent in lungs compared to liver metastases (**Fig. 1D,E** and **Extended Data** Fig. 3A-G). Alongside, compared to lung metastases, the barcode composition of liver metastases exhibited higher similarity to CTCs (**Fig. 1F,G** and **Extended Data** Fig. 3H,I). To test whether indeed lungs and liver provide distinct microenvironments that either support or suppress cancer cell intravasation, we performed three types of injections of GFP-luciferase-labelled or Thy1.2-labelled MVT1 cells in immunodeficient and immunocompetent (FVB, Thy1.1 background) mice, respectively: orthotopic mammary fat pad injection to establish a primary tumor, intravenous (i.v.) injection to obtain lung metastasis and splenic injections to obtain liver metastases. We then analyzed CTC numbers two weeks after cancer cell injection. This relatively early timepoint was chosen to specifically examine intravasation events originating from these target sites before (substantial) further spread to other organs could occur. Strikingly, we observed that the liver environment was strongly favoring CTC intravasation before any of the other sites, in both immunodeficient and immunocompetent models (**Fig. 1H-J**). Together, these results suggest a differential behavior of breast cancer cells as a function of their visceral metastatic site of residence, whereby cells within liver metastases spread further, while cells within lung metastases are less likely to intravasate.

### The liver boosts metastatic properties of breast cancer cells

Next, we set out to prospectively verify whether, beyond differential intravasation rates, cancer cells that metastasized to liver or lungs would harbor a different ability to effectively achieve secondary seeding. We employed orthotopic mammary fat pad injection of two TNBC cell lines, MVT1 and 4T1, both lentivirally labelled with a GFP-luciferase construct. Upon primary tumor growth and spontaneous dissemination to visceral sites, we isolated cancer cells that metastasized to either lungs or liver and transplanted equal numbers (up to 15’000 cells each) through the tail-vein of recipient, tumor-free NSG mice to measure metastatic outgrowth (**Fig. 1K**). We found that liver-resident breast cancer cells gave rise to significantly higher metastatic outgrowth in recipient mice compared to their lung-resident counterpart (**Fig. 1L,M**). Altogether, these results not only suggest that liver-resident breast cancer cells are more likely to spread, but also that the liver microenvironment enhances secondary metastasis-forming features of resident lesions, compared to the lung microenvironment.

### Organ-specific gene expression patterns of breast cancer cells

To gain insights into the differential metastatic propensity of liver-resident vs lung-resident breast cancer cells, we sought to investigate molecular features of both cancer cells and their niche at each site at a single cell resolution. We first transduced GFP-labelled breast cancer cells with a niche labelling system (*19*), resulting in constitutive expression of a soluble mCherry form that is taken up by neighboring cells, enabling prospective niche labeling upon dissemination to lungs and liver. In this context, we isolated cancer (GFP-positive, mCherry-positive) and niche (mCherry-positive) cells from the primary tumor, liver and lungs and performed 10X single cell RNA sequencing (**Fig. 2A**). We first focused on the analysis of 28’105 cancer cells, of which 9’708 derived from the primary tumor, 9’442 derived from liver metastases and 8’955 derived from lung metastases. Unsupervised clustering analysis identified 11 distinct clusters (**Fig. 2B,C**), 6 of them being specific to the primary tumor, lung or liver metastases (**Fig. 2D**). Particularly, we observed clusters 11 and 4 highly enriched within the primary tumor, clusters 8 and 6 characteristic of lung-resident cancer cells, and clusters 7 and 12 specific to liver-resident metastases. Based on this, we next investigated gene markers that characterize these clusters as well as each site, likely to reveal organ-specific gene expression dynamics of breast cancer cells. We adopted a pathway enrichment and gene ontology biological process analysis strategy and pinpointed distinct gene sets upregulated in breast cancer cells resident in each organ (**Fig. 2E**) as well as within each of the gene expression clusters found to be most distinctive of the primary tumor-, lung- and liver-resident cells (**Fig. 2F**). As expected, we found high concordance between these two approaches. Gene sets enriched within the primary tumor included a variety of processes related to translation, including eukaryotic translation initiation and formation of free 40S subunits. In contrast, liver-resident breast cancer cells displayed activity of a variety of metabolic processes related to energy generation, such as glycolysis, metabolism of carbohydrates, citric acid cycle and respiratory electron transport. All these shared a common feature of being related to ATP generation, consistent with a higher metastatic activity of liver-resident cells. On the other side, lung-resident breast cancer cells were enriched in processes such as apoptotic response or detoxification of reactive oxidative species, indicating a higher degree of cellular stress (**Fig. 2E,F** and **Extended Data** Fig. 4, **Supplementary Tables 3,4**). To explore the generality of these findings, we then performed the same analysis on the transcriptome of cancer cells isolated from the primary tumor, lungs and liver metastases obtained from xenografts injected in the mammary fat pad with the patient-derived CTC model BR16 (*20*). Also in this case, we observed enrichment of terms related to translation and protein folding in the primary tumor, energy generating processes such as NAD-related metabolic processes in the liver, and higher levels of cell stress (including apoptosis) in the lung-resident cells (**Extended Data** Fig. 5**, Supplementary Table 5**).

**Fig. 2.**
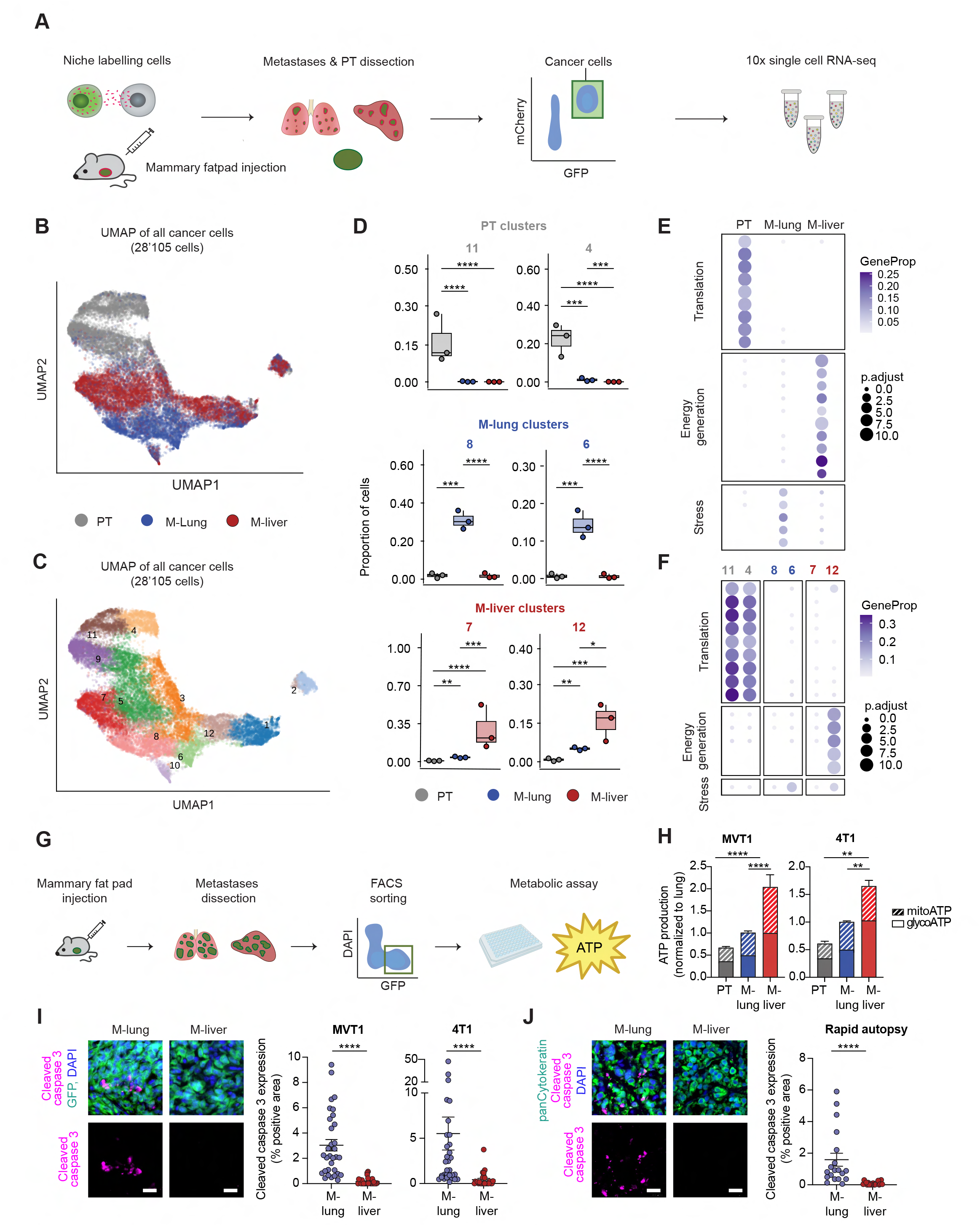
Distinct molecular profiles of liver-resident and lung-resident breast cancer cells. **A.** Overview of the niche-labelling experiment performed with the MVT1-GFP-luciferase model and with a specific focus on the cancer cells component. **B-C.** Uniform Manifold Approximation and Projection (UMAP) embedding colored by **B)** organ of origin and **C)** cell cluster number. **D.** Box-plot representation of specific cell clusters enriched within the primary tumor (PT), liver (M-liver) and lung (M-lung) metastases, along with associated abundances at each anatomical site (propeller derived adjusted *P* value; *, *P* < 0.05; **, *P* < 0.01; ***, *P* < 0.001; ****, *P* < 0.0001). **E-F.** Enriched reactome pathways and gene ontology terms in **E)** organ specific gene expression clusters and in **F)** all cancer cells associated with a given organ. Top 20 pathways for each group with hypergeometric test adjusted *P* < 0.01 are shown. Displayed is a condensed set of ontologies following simplification through semantic similarity. The complete pathway list is provided in Supplementary Tables 3 and 4. **G.** Illustration of the experimental design for the assessment of metabolic properties of breast cancer cells isolated from different sites. **H.** Plot showing the total ATP production rate generated in glycolysis and oxidative phosphorylation obtained from oxygen consumption rate (OCR) and extracellular acidification rate (ECAR). **, *P* < 0.01, ****, *P* < 0.0001 by Mann Whitney test. **I**. Representative images (*left*) of lung (M-lung) and liver (M-liver) tissue sections stained for DAPI (*blue*), GFP (*green*) and cleaved caspase 3 (*magenta*) and dot plot showing the percentage of cleaved caspase 3 positive signal within GFP-positive cancer cells in the MVT1-GFP-luciferase model (*middle*) and the 4T1-GFP-luciferase model (*right*) (*n* = 3 mice for each model). **J**. Representative images (*left*) of lung (M-lung) and liver (M-liver) tissue sections derived from breast cancer patients obtained through rapid autopsy and stained for DAPI (*blue*), pan-cytokeratin (*green*) and cleaved caspase 3 (*magenta*) and dot plot showing the percentage of cleaved caspase 3 positive signal within pan-cytokeratin-positive cancer cells (*right*) (*n* = 3 patients). **I-J** Scale bar, 20 µm; ****, *P* < 0.0001 by Mann Whitney test.

We next followed-up with orthogonal approaches to validate the observed increase in metabolic activity of breast cancer cells when exposed to the liver microenvironment, as well as their increased apoptotic rates when exposed to the lung microenvironment. To this end, we injected two different TNBC cell lines, MVT1 and 4T1, in the mammary fat pad of NSG mice, generating primary grafts with a high propensity to spontaneously metastasize to visceral sites. We then FACS-isolated live cancer cells from the primary tumor, liver and lung metastasis of each model and performed a Seahorse assay to measure the oxygen consumption rates (OCR) and extracellular acidification rates (ECAR) within each site, inferring ATP generation capacity (**Fig. 2G**). In line with our site-specific gene expression analysis, we found that the total ATP generated by TNBC cells within the liver microenvironment was higher than that of primary tumor and lung (**Fig. 2H**). We also confirmed, *via* immunostaining analysis, higher cleaved caspase 3 levels in TNBC cells within the lung microenvironment (**Fig. 2I**). Lastly, we took advantage of the availability of matched liver and lung metastasis specimens of breast cancer patients obtained through a rapid autopsy program performed at the University of Texas Southwestern Medical Center, Dallas. While we were not able to perform a Seahorse assay due to insufficient viability of the cells obtained directly from patient samples overseas, we were able to validate the increase in cleaved caspase 3 within lung metastases by means of immunostaining analysis (**Fig. 2J**).

Together, these results reveal distinct molecular features of breast cancer cells as a function of their visceral organ of residence, suggesting that the tumor microenvironment shapes their biological features in an organ-dependent manner. While breast cancer cells within liver metastases display high metabolic activity and ATP generation capacity mirrored by an enhanced ability to spread further, breast cancer cells within lung metastases appear more subject to stress factors, resulting in higher apoptotic rates and a lower ability to engage in secondary seeding. This raises an intriguing question regarding the identity and function of cells composing the breast cancer metastatic niche in both liver and lungs.

### Interaction between breast cancer cells and their surrounding niche in liver and lungs

We next shifted our focus to the niche component of the sequencing data (**Fig. 3A**), aiming to pinpoint cell types – and their gene expression features – enriched in the tumor microenvironment at different sites. To this end, using the MVT1 model, we isolated, sequenced, and analyzed 12’558 niche cells, of which 4’257 microenvironment cells located within the primary tumor, 3’854 niche cells within lung metastases, and 4’447 niche cells within liver metastases. In parallel, we performed single cell 10X RNA-sequencing of 5’557 lung and 4’740 liver cells obtained from healthy tumor-free mice, enabling a better understanding of baseline, organ-intrinsic features. First, we found substantial differences in the composition of the niche at different sites. While elevated myeloid cell levels are observed in the metastatic niches as compared to healthy lung and liver tissues, we found a marked enrichment of macrophages within the microenvironment of the primary tumor and lungs, while they represent a relatively smaller fraction of the liver metastatic niche (93% in primary tumor, 85% in lungs, 29% in liver; **Fig. 3B,C** and **Extended Data** Fig. 6). When comparing gene expression profiles of specific cell types of the metastatic niche at different locations, we find site-associated features (**Extended Data** Fig. 7), suggesting for instance the presence of alveolar macrophages within the lung metastatic niche and Kupfer cells and infiltrating macrophages within the liver metastatic niche (**Extended Data** Fig. 7,8). Among the most striking differences, however, we found the liver metastatic niche to be unique in respect to other locations, displaying a remarkable enrichment in endothelial cells, consistently with a high representation of this cell type in the healthy liver (**Fig. 3 B,C** and **Extended Data** Fig. 6). This suggests a possible involvement of the endothelium in shaping metastasis-forming features of liver-resident breast cancer cells.

**Fig. 3.**
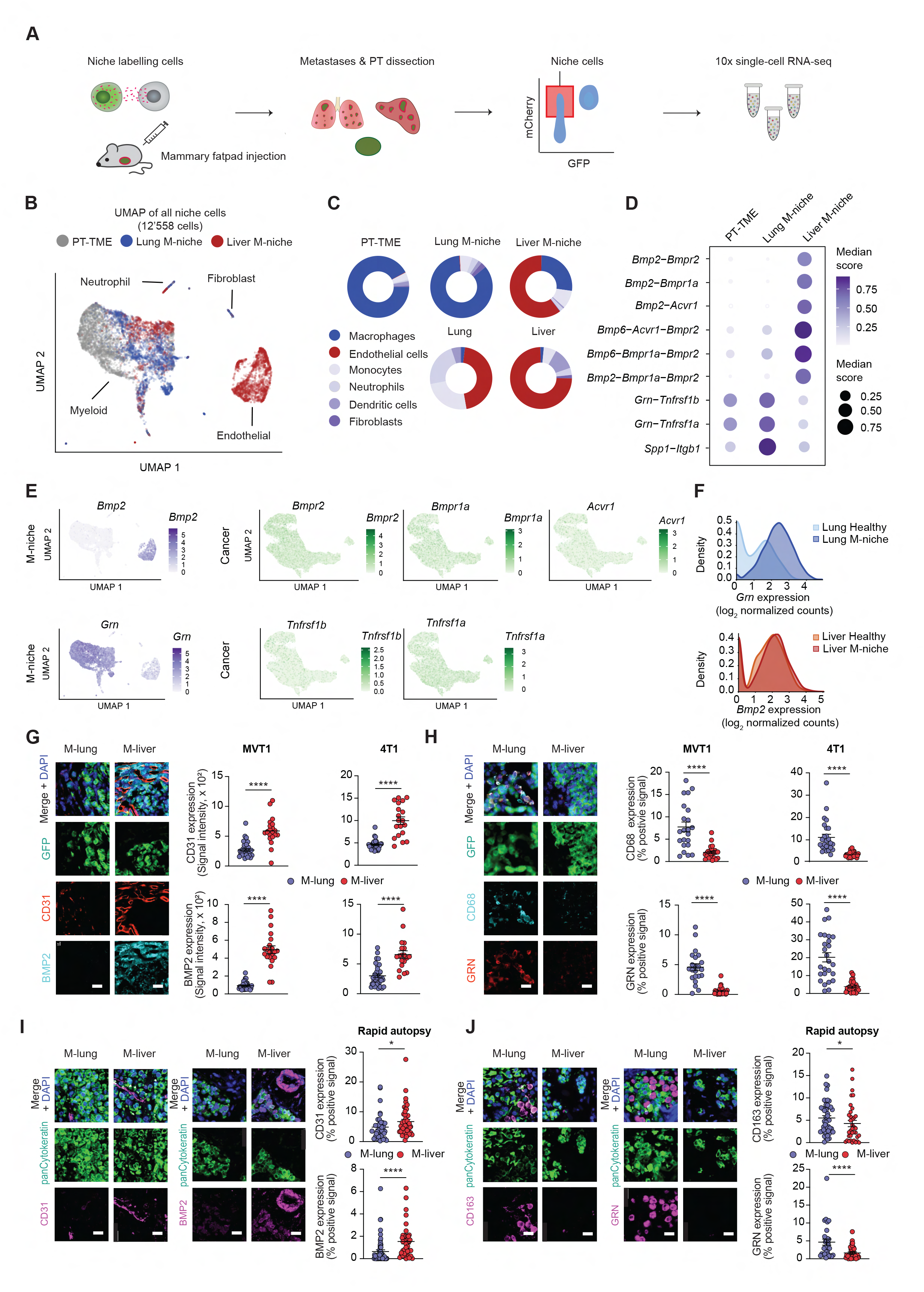
Metastatic niche composition of liver and lungs in breast cancer. **A.** Overview of the niche-labelling experiment with the MVT1-GFP-luciferase model with a specific focus on metastatic niche cells. **B.** Uniform Manifold Approximation and Projection (UMAP) embedding of the primary tumor microenvironment (PT-TME) and metastatic niche (M-niche) cells colored by organ. Annotations indicate the locations of estimated cell types within UMAP. **C.** Pie charts displaying cell type distributions within the tissue microenvironment of the primary tumor (PT-TME) and the metastatic niches (M-niche) of lungs and liver, as well as the composition of healthy lung and liver tissues. **D.** Dot plot representing selected interactions between cytokine ligands (niche cells) and their receptors (cancer cells) at each site. Dot color and size correspond to the average interaction score scaled for each ligand-receptor pair using minimum-maximum normalization. **E.** UMAP displaying cytokine and receptor expression in niche cells and cancer cells, respectively (refer to panel **B** and Fig. 2B for the cell type and organ of origin). **F.** Plot showing the expression levels of *Grn* in macrophages of the healthy and metastatic lungs (*top*), as well as the expression levels of *Bmp2* in endothelial cells of the healthy and metastatic liver (*bottom*). **G**. Representative images (*left*) of lung (M-lung) and liver (M-liver) tissue sections stained for DAPI (*blue*), GFP (*green*), CD31 (*red)* and BMP2 (*cyan*) and dot plots showing the intensity of the respective immunofluorescence signals in MVT1-GFP-luciferase (*middle*) and 4T1-GFP-luciferase mouse models (*right*) (*n* = 3 mice for each model). **H**. Representative images (*left*) of lung and liver tissue sections stained for DAPI (*blue*), GFP (*green*), CD68 (*cyan)* and GRN (*red*) and dot plots showing the percentage of threshold-set positive signal of the respective immunofluorescence signals in MVT1-GFP-luciferase (*middle*) and 4T1-GFP-luciferase mouse models (*right*) (*n* = 3 mice for each model). **I**. Representative images (*left, middle*) of lung and liver metastasis tissue sections stained for DAPI (*blue*), pan-cytokeratin (*green*), CD31 (*magenta*) and BMP2 (*magenta*) and dot plots showing the percentage of threshold-set positive signal of the respective immunofluorescence staining (*right*) in breast cancer patients (*n* = 3 patients). **J**. Representative images (*left, middle*) of patient lung and liver metastasis tissue sections stained for DAPI (*blue*), pan-cytokeratin (*green*), CD163 (*magenta)* and GRN (*magenta)* and dot plots showing the percentage of threshold-set positive signal of the respective immunofluorescence staining (*right*) in breast cancer patients (*n* = 3 patients). Scale bar, 20 µm. *, *P* < 0.05, ****, *P* < 0.0001, by Mann Whitney test.

By narrowing down our focus to macrophages and endothelial cells as the key components of the lung and liver metastatic niches, respectively, we identified gene expression alterations in the metastatic niche as compared to the normal healthy tissue (**Extended Data** Fig. 9A,B). Interestingly, the macrophages of the lung metastatic niche showed an enrichment of processes related to cell cycle and proliferation as compared to their counterpart in normal lungs, coinciding with the prominent abundance (and likely, activation) of this population in the lung metastatic niche. Macrophages within the liver metastatic niche, conversely, were characterized by an enhancement of the pathways related to the energy generating metabolic pathways and translation (**Extended Data** Fig. 9C). On the other side, the endothelial cells in the liver metastatic niche showed an upregulation of pathways related to translation, migration and activation compared to endothelial cells within the healthy liver (**Extended Data** Fig. 9D).

We next sought to define key interactions between the cancer cells and different components of their metastatic niche, focusing on surface receptors present on cancer cells and corresponding ligands secreted by the niche counterpart, based on known and recognized interaction pairs (*21*, *22*). For each ligand-receptor pair, we scored interactions using the product of average ligand expression in niche cells and the average receptor expression across all cancer cells. We selected interactions of significant score (*P* < 0.05) and then identified those that were present preferentially in liver or lungs (**Fig. 3D** and **Extended Data** Fig. 10A-C), with a ratio of at least 2-fold increase over the other counterpart. This enabled us to identify *Grn* and *Ssp1* secreted by macrophages and mirrored by their receptors *Tnfrsf1a/b* and *Itgb1*, respectively, expressed by cancer cells within lung metastases. On the other hand, for liver metastases we identified interactions between endothelial *Bmp2* and *Bmp6* cytokines and *Bmpr2 / Bmpr1a* / *Acvr1* receptors on cancer cells (**Fig. 3D, Extended Data** Fig. 10A-C). The expression of these ligands and receptors in the niche and in cancer cells are depicted in **Fig. 3E** and **Extended Data** Fig. 11, showcasing a highly specific presence of *Bmp2* and *Bmp6* cytokines in liver endothelial cells, as well as *Grn and Ssp1* expression in lung macrophages, with *Bmp2* and *Grn* displaying the highest expression in niche-cells of liver and lung metastatic niches, respectively (**Fig. 3E** and **Extended Data** Figs. 10C, 1**1**). Of note, while *Grn* expression levels are augmented in the macrophages of the lung metastatic niche when compared to their counterpart in healthy lungs, *Bmp2* is expressed at comparable levels by the endothelial cells of both in the healthy and metastatic livers (**Fig. 3F**).

Focusing on the two predominant niche cell types and the respective cytokines, we set out to validate, using orthogonal approaches, the macrophage and endothelial cell content as well as the respective GRN and BMP2 levels in lung and liver metastases of mouse models and patient samples. With immunostaining analysis of mouse samples from 4T1- and MVT1-GFP-luciferase orthotopic models, we found higher levels of CD31-positive cells in the liver metastatic niche, indicating a larger extent of vascularization and a simultaneous enrichment in BMP2 expression (**Fig. 3G**). At the same time, the CD68-positive macrophage content, including GRN-expressing cells, was enhanced in the lung tissue of these models (**Fig. 3H**), in line with our single-cell RNA sequencing data. To assess the relevance of our findings in a clinical context, we again made use of patient-matched liver and lung metastases samples obtained through rapid autopsy. Consistently, we found that the liver metastatic niche displayed a higher level of vascularization and showed higher expression of BMP2 (**Fig 3I**), and that CD163-positive macrophages were more abundant in the lung metastatic niche, along with a higher GRN expression (**Fig. 3J**).

Altogether, these results highlight a distinct composition of the metastatic niche within lung and liver in breast cancer mouse models and patients. While lung metastatic cells mainly interact with GRN-secreting alveolar macrophages, liver metastatic cells appear to rely on BMP2-secreting endothelial cells. Given the enhanced metastatic ability of liver-resident breast cancer cells, we next focused on validating the functional relevance of BMP2 signaling in the spread of breast cancer.

### BMP2 promotes metastasis-forming properties of breast cancer cells

Given its potential targetability, we next sought to validate our findings on BMP2, particularly by testing its ability to modulate functional features and metastatic potential of breast cancer cells. To this end, we first stimulated MVT1 cells with BMP2 at 100 ng/ml for 24 hours, and simultaneously knocked out *Bmpr2*, *Bmpr1a* and *Acvr1* receptors that mediate BMP2 signaling, to test whether this would abrogate BMP2-induced effects (*23–25*). Given the pronounced metabolic activity of liver-resident breast cancer cells, we tested whether BMP2 stimulation had an effect on ATP generation (**Fig. 4A**). Upon confirming that BMP2 treatment resulted in increased levels of phospho-SMAD1/5 (*26*) (**Extended Data** Fig. 12), we found that BMP2 stimulation increased ATP production in MVT1 cells, consistently with previous *in vivo* results, and that this effect was abolished by means of BMP2 receptors knockout (**Fig. 4B**).

**Fig. 4.**
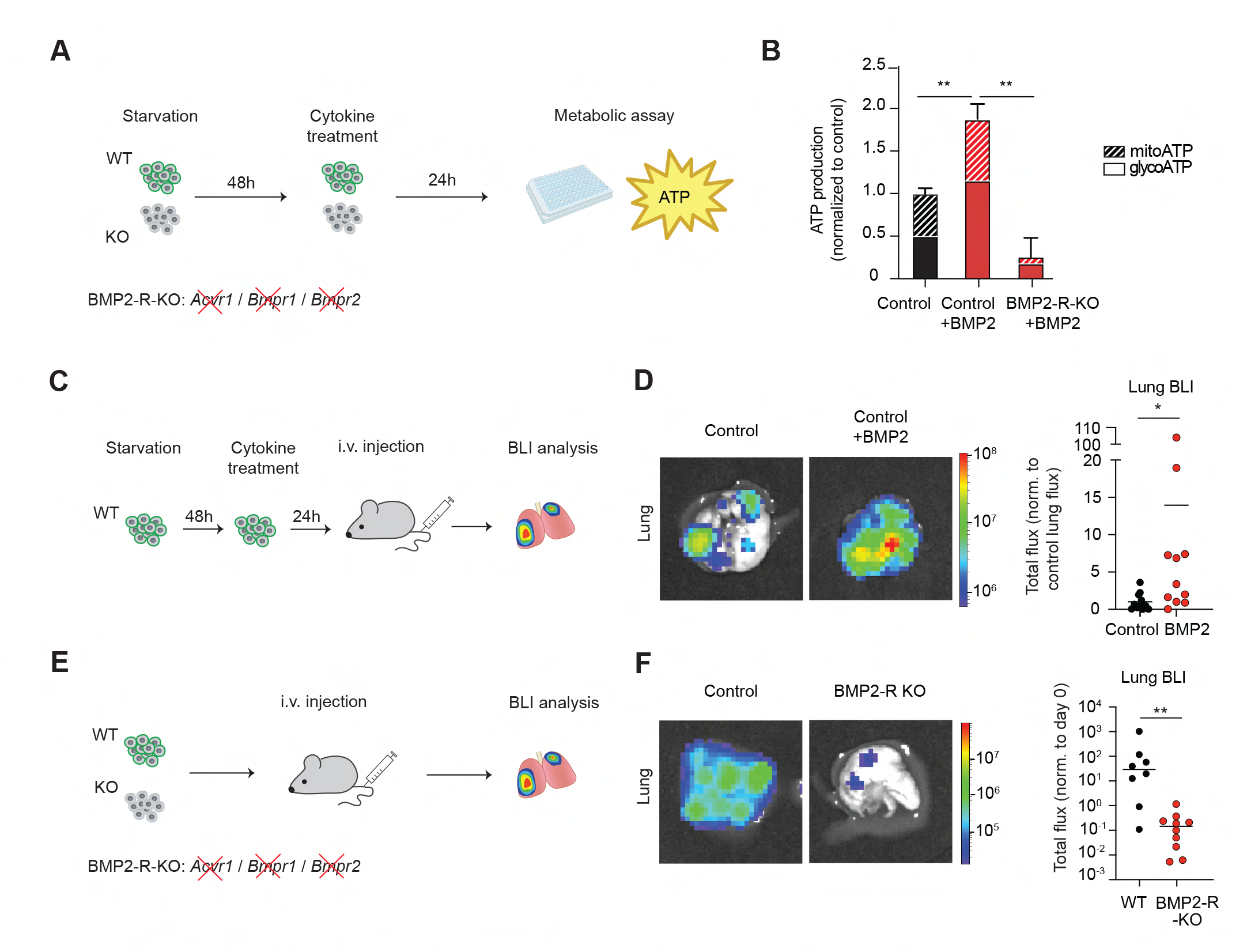
BMP2 axis targeting reduces breast cancer metastasis. **A.** Illustration of the seahorse assay using untreated (control) and BMP2 treated MVT1 GFP-luciferase cells, with or without *Acvr1*-*Bmpr1*-*Bmpr2* receptors knockout (BMP2-R KO). **B.** Plot showing the total ATP production rate generated in glycolysis and oxidative phosphorylation obtained from oxygen consumption rate (OCR) and extracellular acidification rate (ECAR) for untreated (control) and BMP2 treated MVT1-GFP-luciferase cells, with or without *Acvr1*-*Bmpr1*-*Bmpr2* receptors knockout (BMP2-R KO). **, *P* < 0.01 by Mann-Whitney test. **C.** Experimental design to explore the effect of BMP2-stimulation on secondary seeding capacity of MVT1-GFP-luciferase cells. **D.** Representative images (*left*) and quantification (*right*) of bioluminescence signal of lungs in NSG mice injected with MVT1-GFP-luciferase cells untreated (control) and treated with BMP2. *, *P* < 0.05 by Mann-Whitney test. **E.** Experimental design to explore the role of BMP2-signalling on secondary seeding capacity of MVT1-GFP-luciferase control cells or cells carrying *Acvr1*-*Bmpr1*-*Bmpr2* receptors knockout (BMP2-R KO). **F.** Representative images (*left*) and quantification (*right*) of bioluminescence signals of lungs of NSG mice injected with control and BMP2-R KO MVT1-GFP-luciferase cells. **, *P* < 0.01 by Mann-Whitney test. WT, wild-type; BLI, bioluminescence imaging.

Based on this finding, to mimic the effect of a BMP2-rich microenvironment as found in liver metastases, we pretreated MVT1-GFP-luciferase cells *in vitro* with 100ng/ml of BMP2 for 24 hours, before intravenously injecting them into tumor-free recipient NSG mice (**Fig. 4C**). With the purpose to assess the impact of this treatment on secondary metastatic seeding, we prospectively measured new metastasis formation in the lungs. In line with previous findings, treatment with BMP2 led to an enhanced metastatic ability of MVT1 cells compared to untreated control cells (**Fig. 4D**). Further, to investigate whether the abrogation of BMP2-mediated signaling could have the opposite effect, we measured lung metastasis – as a proxy for secondary seeding capacity – upon intravenous injection of MVT1 control cells or cells carrying *Bmpr2*, *Bmpr1a* and *Acvr1* triple-receptor knockouts (**Fig. 4E**). Consistently, we found that disruption of BMP2 signaling impaired the metastatic capacity of MVT1 cells, as demonstrated by bioluminescence imaging (**Fig. 4F**).

Altogether, these findings highlight a prominent role of BMP2 signaling in boosting metastasis-forming properties of breast cancer cells that reside within the liver metastatic niche. Given its pivotal role in metastasis, targeting BMP2 signaling, either by antagonizing BMP2 itself or blocking its receptors at the level of cancer cells, may represent a promising therapeutic strategy for limiting secondary spread from liver metastases of breast cancer.

## Discussion

A better understanding of the role of the niche cells within different metastatic sites in shaping the properties of newly resident cancer cells is needed to better understand cancer cell biology as a function of their location. This will shed light on processes that occur relatively late during a patient’s history, yet, may have a broad relevance for the development of new therapeutic options to be tested in the context of a metastatic disease. Previously, it has been suggested that the bone metastatic niche may invigorate resident cancer cells to further disseminate (*26*), a finding that is highly relevant for hormone-driven cancers. Here, we provide first insights into the metastatic niche of lung and liver metastases of breast cancer. We observe differences in cellular composition, as well as in the cytokine crosstalk between cancer and niche cells within these exclusive sites, impacting on patterns of disease progression, metastasis-relevant characteristics, as well as disease outcome.

Our findings suggest that the liver microenvironment, mainly *via* the action of BMP2-secreting endothelial cells, invigorates resident breast cancer cells and promotes secondary spread in the form of metastasis-to-metastasis dissemination. This is in line with a higher ability of liver metastases to generate CTCs as well as new metastatic lesions, compared to breast cancer cells that are resident in the lung. For clarity, while lung metastases of breast cancer appear to be subjected to higher stress/apoptosis levels compared to their liver-resident counterpart and display a poorer ability to further metastasize, by no means our data should imply that lung metastases are not disease-relevant. In contrast, it should be clear that breast cancer cells within the lung microenvironment are still well-capable to proliferate and colonize, yet in a more localized fashion, leading to disease progression.

We envision several scenarios that could benefit from these findings. For instance, on a therapeutic angle, TNBC patients with oligometastatic disease could be treated differently as a function of disease location, whereby patients with metastases localized exclusively to the liver could profit from BMP2 targeting alongside standard-of-care treatments, aiming to interfere with disease progression.

Although our study is mostly relevant for TNBC given its marked propensity to metastasize to visceral sites, several other cancer types do spread to liver and lungs. The question of whether our findings on the niche composition on these sites as well as the changes that occur in liver-resident vs lung-resident breast cancer cells also occur in other cancer types with similar metastatic patterns remains unanswered. Uncovering this unexpected dichotomy for sites that have been associated with TNBC should stimulate further investigations in this context, aimed to better map metastatic behaviors, biology and vulnerabilities not only from a generic cancer cell perspective, but also from a geographical standpoint.

## Methods

### Patient data collection and survival analysis

Patient history was collected in line with the principles of the Declaration of Helsinki. Ethical approval was obtained by the local Ethical Committee of Nordwest-und Zentralschweiz (EKNZ, BASEC 2020-03023) and the Kantonale Ethikkommission Zürich (BASEC 2023-00791). Briefly, in this retrospective longitudinal study, we analyzed data from 318 metastatic (Stage IV) breast cancer patients treated between 1972 and 2019 in two Swiss hospitals, the University Hospital Basel and the University Hospital Zürich. For each patient we collected demographic characteristics (age), date of death, primary tumor information (date of diagnosis, stage, histology, immunohistochemical characteristics), and metastasis diagnosis (including anatomical location and date) to reconstruct the sequence of metastatic events. Mono-metastasis as a first event was defined as metastasis diagnosed in a single organ concurrently or upon primary diagnosis. We defined overall survival (OS) as the time between the diagnosis of the first metastatic event and the death from any cause, and time to next event as the time between the first metastasis and the next metastasis or death.

The Surveillance, Epidemiology, and End Results (SEER) Program database (SEER*Stat version 8.4.4) was utilized to case listing of the SEER research data from 17 registries, based on the November 2021 submission, covering the years 2000-2019. For this analysis, we selected cases of female malignant breast cancer with the subtype ‘HR-/HER2-’, as well as cases of colon and rectal cancer, and pancreatic cancer, all with metastasis at diagnosis and with at least one subsequent record.

### Statistical analysis of patient data

Survival curves describing time to next event and OS were estimated using Kaplan-Meier Survival analysis. Survival curves of the different metastatic sites were compared using long-rank test. Hazard ratios and associated 95% confidence intervals were derived from a Cox proportional hazard model with metastatic location as the only independent variable.

### Patient samples

Patient-derived matched liver and lung metastasis samples from breast cancer patients used for immunostaining analysis were obtained from a rapid autopsy program carried out at the University of Texas Southwestern Medical Center, Dallas, TX, USA. Cancer patients were enrolled under the approved protocol #STU-2020-0628 from November 17, 2021, to November 7, 2023, USA. Following the autopsy, the samples were immediately fixed in 4% PFA and subsequently prepared for paraffin embedding.

### Cell lines

The MVT1mouse mammary carcinoma cell line was a kindly provided by Mohamed Bentires-Alj (University of Basel) and originally obtained from a collection of immunocompetent mouse models collected by the Lalage M. Wakefield (*28*). The 4T1 mouse mammary carcinoma cell line was obtained from the American Type Culture Collection (ATCC #CRL-2539). The cell lines were grown in Dulbecco’s modified Eagle medium (DMEM, Gibco, #11330-057) supplemented with 10% fetal bovine serum (Gibco, 10500064) and antibiotic/antimycotic (Gibco, #15240062) in a humidified incubator at 37°C with 20% O_2_ and 5% CO_2_. For *in vivo* experiments, the cells were transduced with lentiviruses carrying GFP-luciferase. Human breast CTC-derived BR16 cells were generated as previously described (*29*) and propagated as suspension cultures in a humidified incubator at 37 °C with 5% O_2_ and 5% CO_2_.The cell lines were confirmed negative for mycoplasma contamination.

### Mouse experiments

All mouse experiments were carried out according to institutional and cantonal guidelines (approved mouse protocol number 3053, cantonal veterinary office of Basel-City; and approved mouse protocol number 33688, cantonal veterinary office of Zurich). Experimental endpoints that were allowed in our approved license included tumor-related parameters such as a maximum tumor size of 2,800 mm^3^ or moderate ulceration, as well as appearance and behavior parameters such as hunching, piloerection or decreased activity. These limits were not exceeded in any of the experiments. Experimental setups and calculations were in accordance with the 3R principles. All mice were randomized before the start of each experiment, but blinding was not carried out. NSG (NOD-scid-Il2rgnull; The Jackson Laboratory, #005557) female mice were kept in pathogen-free conditions, according to institutional guidelines. Animals were kept in a standard light-cycle photoperiod (12 h light/12 h dark).

### Orthotopic mammary fat pad, tail vein and splenic injections and CTC isolation

Orthotopic mammary fat pad cancer lesions were generated in 10-12-week-old NSG females by injection with 100’000 MVT1-GFP-luciferase, MVT1-Thy1.2, 4T1-GFP-luciferase or BR16-GFP-luciferase breast cancer cells in 50 μl of 50% Cultrex PathClear Reduced Growth Factor Basement Membrane Extract (R&D Biosystems, #3533-010-02) in phosphate-buffered saline (PBS). For intravenous injections, 10’000-100’000 MVT1-GFP-luciferase, MVT1-Thy1.2, and 4T1-GFP-luciferase cells were prepared in a maximum volume of 50 µl and injected into the tail vein using an insulin syringe. For liver-targeting injections, mice were anesthetized, the spleen exposed *via* lateral abdominal incision and 100’000 MVT1-GFP-luciferase or MVT1-Thy1.2 cells in 50 μl PBS were injected directly into the spleen. For metastasis recording and monitoring, mice were imaged with IVIS bioluminescence (PerkinElmer, Living image software) under anesthesia. At the experiment termination, the blood was drawn for CTC analysis (cellular barcoding and intravasation rate experiments) and organs were dissected (all experiments). Blood samples were processed with the Parsortix Cell Separation System (ANGLE) and Cell Separation cassettes (GEN3D6.5), from which the captured CTCs (GFP-positive, CD45-negative) were subsequently released.

### Fluorescence-Activated Cell Sorting (FACS)

Single cell suspensions derived from mouse tissues were obtained by Collagenase IV (Sigma, #C5138-1G) digestion (1 mg/ml) at 37°C for 2×10 minutes, with pipetting in between to disrupt cell clusters, subsequent addition of EDTA (5 mM) and wash in 2% FBS in PBS to quench the reaction. The cell suspensions were filtered through a 40-μm cell strainer (BD Bioscience) and red blood cells were lysed with ACK Lysing Buffer (Thermofisher, #A1049201). Equal numbers of GFP-positive live cells were sorted from lungs and liver. Single cancer cells were selected following doublet discrimination and DAPI dye exclusion. For each sample, a post-sort quality and purity control was performed to ensure that >95% of sorted cells were confirmed to be GFP-positive and DAPI-negative.

### Direct metastatic potential assay

To assess the metastatic potential of lung- and liver-resident cells we first performed orthotopic mammary fat pad injection of MVT1-GFP-luciferase or 4T1-GFP-luciferase cells. Following tumor development, lungs and livers were collected at terminal time-point, upon which metastases were dissected and FACS sorted. Equal numbers of lung- and liver-resident cells (up to 15’000) were injected into tail vein of recipient NSG mice. Metastasis onset and growth rate in lungs were non-invasively monitored on a weekly basis with the IVIS bioluminescence (BLI) system. The experiment was terminated 10 days post-injection and metastasis to lungs and liver were assessed with BLI. For the assessment of metastatic potential of cytokine-treated cells (see section below), 10’000 MVT1-GFP-luciferase cells were injected intravenously into recipient mice and the metastatic burden was assessed *ex vivo* through IVIS BLI imaging 10 days post-injection.

### Cellular barcoding

MVT1-GFP-luciferase cells were transduced with a barcode lentiviral library from Cellecta (CloneTracker XP™ 5M Barcode-3’ Library in pScribe4M-RFP-Puro) at MOI<0.1 to maximize the likelihood of a unique barcode integration in each individual cell. Barcode-transduced GFP-positive and RFP double-positive cells were FACS-sorted, both for pre-injection barcode pool controls and for orthotopic injection into mice (100’000 cells per mouse). At the end of the experiment, we analyzed all organs for metastases by IVIS BLI imaging and collected metastatic deposits, lysed them and extracted their DNA. Barcode regions were amplified with a set of staggered primers flanking the barcode locus, followed by a nested PCR using primers that introduce i5 and i7 adapters to bind to the sequencing flow cell and a binding site for the sequencing primer. Samples were sequenced with Illumina Novaseq 6000, using 150bp Paired End. Primer sequences targeting barcodes are reported below:

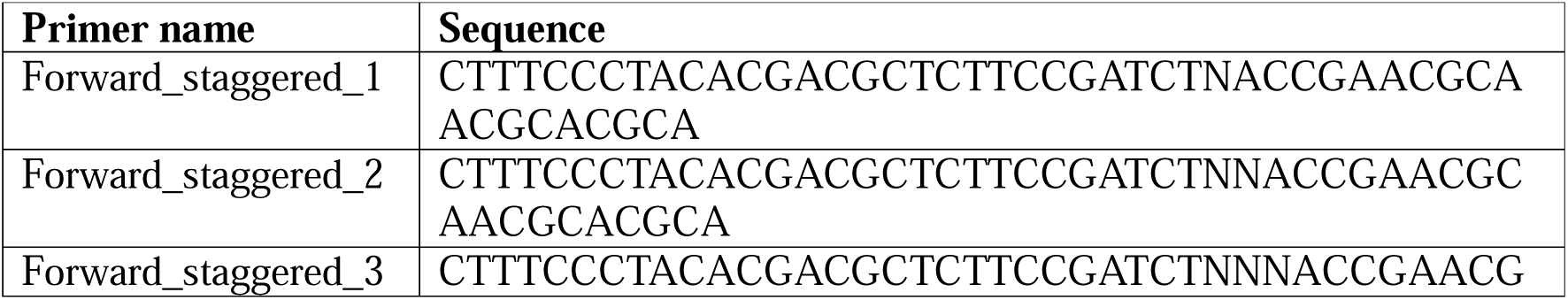
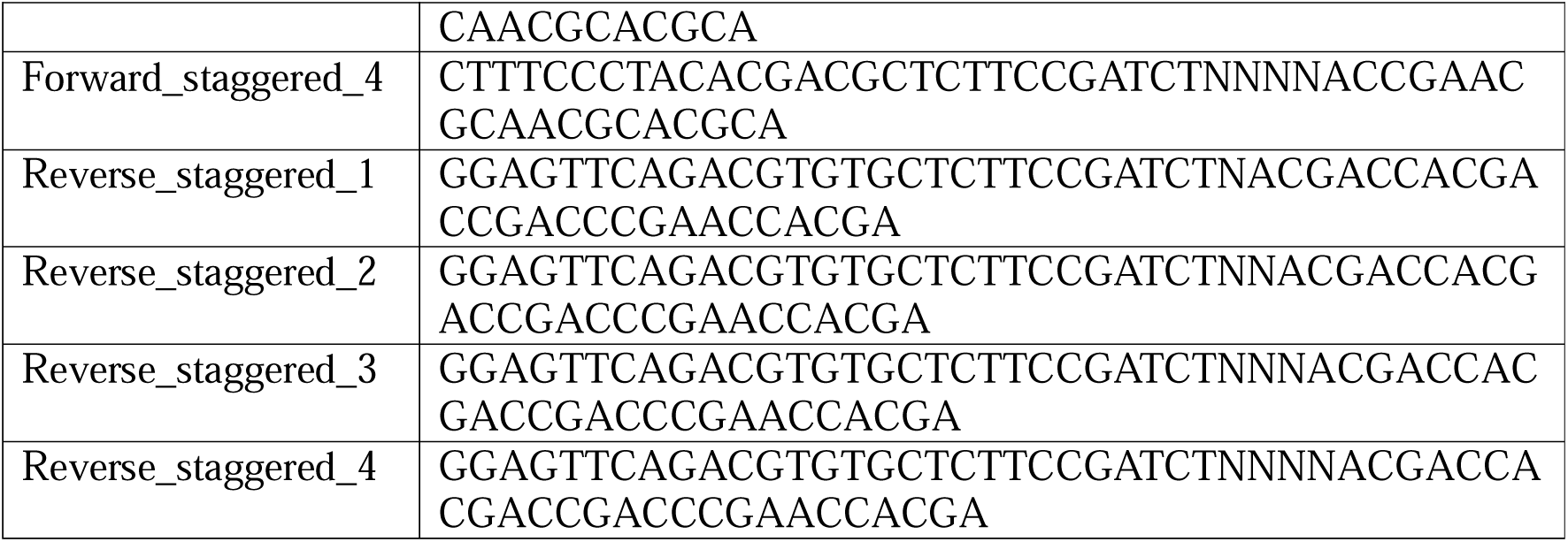

### Barcode sequence analysis

Sequencing reads were processed using cutadapt (v4.2) (*30*), resulting in trimmed reads of lengths 30 bp and 14 bp. The trimmed reads were then aligned to a reference barcode library supplied by Cellecta using Bowtie2 (v2.3.5.1) (*31*) and filtered using Samtools (v1.12) (*32*), retaining only those with a mapping quality score greater than 20. After filtration, reads common to both the 14 bp and 30 bp sets were identified and extracted using Samtools (v1.12) (*32*) and Picard (v2.25.7) (Picard Toolkit. 2019. Broad Institute, GitHub Repository. https://broadinstitute.github.io/picard). The frequency of common reads was then counted for every sample. Barcode threshold of a combined 5 counts per million (CPM) per organ was applied to qualify them as organ-present barcodes. Subsequently, differential analyses were performed using Limma (v3.58.1) (*33*) under standard settings. Barcodes were considered unique for a given organ based on their expression in the organ (CPM>5) and absence-criteria that included an adjusted *P* value threshold below 0.05, an absolute log2 fold change higher than 0.5, a CPM below 0.05% and a scale value below 0.2 in all the remaining samples. Euclidean distances measuring the dissimilarity in barcode expression between lung or liver metastases and CTCs was obtained by analysis of barcodes that exist exclusively in lung and/or circulating tumor cell (CTC) samples when calculating the distance between lung and CTC samples; exclusively in liver and/or CTC samples for the liver and CTC distance.

### Seahorse – Mitochondrial Stress Test

Oxygen consumption was measured using the Seahorse Xfe96 Analyzer (Agilent) according to the manufacturer’s protocol. Cultivated cells or FACS-sorted tumor cells were transferred to a poly-D-lysine–coated 96 well plate and incubated for at least 1 hour in a CO_2_-free incubator in either DMEM or RPMI containing no phenol red with10mM glucose, 1mM pyruvate, and 2mM glutamine at 37°C. A Seahorse Xfe96 Analyzer was used to inject 10 μM oligomycin, 15 μM Carbonyl cyanide-p-trifluoromethoxyphenylhydrazone, and rotenone 1 μM + antimycin A 10 μM. The Seahorse Xfe96 Analyzer was used to measure the oxygen consumption rate (OCR) and extracellular acidification rate (ECAR) at baseline and after each injection. The ATP generated in glycolysis and oxidative phosphorylation was obtained with Seahorse analytics software.

### Immunostaining analysis of mouse and patient samples

Mouse lung and liver tissues were fixed in 4% paraformaldehyde at 4°C overnight, washed in PBS, incubated in 30% sucrose, embedded in OCT and sectioned with cryostat at 10 μm thickness. The sections were then washed in PBS, briefly immersed in 0.01% Triton X-100 in PBS and incubated in 2% donkey serum, 0.01% Triton X-100 in PBS for 1hour. Subsequently the sections were incubated with the primary antibodies in 2% donkey serum, 0.01%Triton-X-100 in PBS overnight (1:250 Cleaved Caspase 3 from Cell Signaling Technology #9661; 1:100 CD31 from Biotechne #AF3628; 1:50 BMP2 from Abcam #214821; 1:200 CD68 from Biorad #MCA1957T; 1:100 Granulin from Biotechne #AF2557), washed three times for ten minutes the next day, and then incubated with the secondary antibodies (1:500 donkey anti-rabbit AF647 from Thermofisher #A-31573; 1:500 donkey anti-goat AF568 from Thermofisher #A-11057; 1:500 goat anti-rat APC from Thermofisher #A-10540; 1:500 donkey anti-sheep AF555 from Thermofisher #A-21436, 1:500) and DAPI in 2% donkey serum, 0.01%Triton-100X in PBS for three hours, washed and mounted.

Patient FFPE samples were sectioned with a microtome at 5 μm thickness, following a standard deparaffinization/antigen retrieval protocol. Next, samples were permeabilized in 5% Triton X-100 in PBS for 20 mins, washed and incubated in 10% donkey serum, 1% BSA, 0.5% Triton X-100 in PBS, subsequently incubated in primary antibodies in 3% donkey serum, 1% BSA, 0.5% Triton X-100 in PBS overnight (1:100 pan-cytokeratin from GeneTex #GTX27753; 1:250 Cleaved Caspase 3 from Cell Signaling Technology #9661; 1:50 CD31 from Abcam ab28364; 1:25 BMP2 from Abcam #14933; 1:200 CD163 from Leica Biosystems #NCL-L-CD163; 1:50 Granulin from Biotechne #AF2420), washed 3 times for 10 minutes the next day, and incubated with secondary antibodies (donkey anti-rabbit AF647 from Thermofisher #A-31573; 1:200 donkey anti-goat AF647 from Thermofisher #A-32849; 1:200 goat anti-mouse AF647 from Abcam #ab150119; 1:200 donkey anti-mouse AF555 from Thermofisher #A-31570) and DAPI in 0.01% Triton X-100 in 3% donkey serum, 1% BSA, 0.5% Triton X-100 in PBS for 3 hours, washed and mounted.

Immunofluorescence imaging was carried out on a Nikon widefield microscope, and images were taken using the 20X lens. All images were analyzed by the Fiji image processing software.

### 10x RNA-sequencing

The niche-labelling cells were obtained by lentiviral transduction of MVT1-GFP-luciferace cells with the sLP-mCherry construct (*19*). 100’000 transduced cells were injected orthotopically into the mammary fat pad of NSG mice. At the end of experiment, we analyzed all organs for metastases and collected metastatic deposits. Cells were FACS-sorted and stained with Hashing antibodies (TotalSeq™-B0301-307 anti-mouse Hashtag 1-7; M1/42; 30-F11PG from Biolegend, #155831, #155833, #155835, #155837, #155839, #155841, #155843) and subsequently pooled to include cancer cells from each mouse in one sequencing flowcell, as well as niche cells from each mouse in one sequencing flowcell to eliminate batch-effect when comparing gene expression across sites for the same biological replicate. Library preparation was performed using Chromium Next GEM single cell 3‘reagent kits v3.1 according to the manufacturer’s instructions (10X Genomics, CA, USA). Samples were sequenced with Illumina Novaseq 6000.

### CRISPR Cas9 knockout generation

MVT1-GFP-luciferase cells were transduced with Cas9-RFP lentiviruses (Sigma-Aldrich, #CAS9RFPV-200UL). Three Cas9-bearing cell clones were isolated and expanded based on their phenotypic resemblance to the parental cell line, then combined into a pool of clones. Subsequently, MVT1-GFP-luciferase-Cas9 cells were transduced with lentiviruses bearing sgRNAs within a pLCKO2 plasmid. Subsequently, knockout clones were obtained and validated by detecting the deletion with DNA sequencing of the PCR products containing the deletion/indel site and comparing the putative CRISPR-Cas9 modified DNA with the DNA of the wild-type, non-targeted cells. We targeted *Bmpr2, Bmpr1a* and *Acvr1* simultaneously at high MOI to maximize the likelihood of multiple knockouts in target cells.

### *In vitro* cytokine treatment

100’000 MVT1-GFP cells per well were seeded in a six-well plate and cultured in growth medium overnight. The following morning, cells were washed three times with PBS and given starvation medium (0.1% FBS in DMEM). After 48 h, the medium was supplemented with BMP2 (R&D Systems, #355-BM-010/CF) at 100ng/ml as previously reported (*34*, *35*). Cells were stimulated for 24 h and then collected upon trypsinization, enumerated using automatic cell counter and 10,000 cells were injected intravenously into 12-week-old female NSG mice.

### Single cell RNA-seq data pre-processing

FASTQ files were demultiplexed and reads were aligned to an extended mouse genome reference (GRCm39 and GENCODE release M29) containing *GFP* and *mCherry* sequences using CellRanger (v.6.1.2, 10x Genomics). FASTQ files from BR16 model were aligned to human genome reference (GRCh18 and GENCODE release 44). Next, we excluded low-quality cells based on the total UMI counts, the number of detected genes and the proportion of mitochondrial reads. Filtering criteria were established for each sequencing batch using adaptive thresholds defined as 3 median absolute deviation in the problematic direction. HTOs were demultiplexed using the HTODemux function from Seurat (v 4.3.0) and cells without an expected HTO or doublets with two HTO assigned were removed from the analysis. Data from different batches were integrated adjusting by library size and correcting batch effects using mutual nearest neighbors (MNNs) method implemented in the fastMNN function R/Bioconductor package batchelor (v1.14.1). After integration, we clustered the data using walktrap method based on principal components of corrected data and visualized using UMAP projection. Niche cells clustering together with cancer cells were removed from the analysis as they might represent potential sorting errors. For the specific analysis of cancer cells (Fig. 2) and niche cells (Fig. 3), and normal organs, split data was integrated, batch corrected and clustered as detailed above. An additional doublet detection to removed doublets containing the same HTO was performed using scDblFinder R/Bioconductor package (v1.13.8).

### Single cell RNA-seq data analysis

We analyzed the differential abundance of cells from different anatomical sites within each cluster (Fig. 2D) using the propeller function from the speckle R package (**v0.0.3** (*36*). Markers for each cluster and anatomical site were detected using scoreMarkers function from R/Bioconductor package scran (v1.26.2) blocking by batch. Genes with a mean AUC≥0.25 in each cluster were used as an input to the over-enrichment analysis (clusterProfiler R/Bioconductor package, v4.6.0) using the Reactome pathways and gene ontology biological processes included in the MSigDB (v2022.1.Mm and v2023.1.Hs). Pathways with an adjusted *P* value < 0.01 were clustered using hierarchical clustering on pairwise semantic similarity based on Jaccard index. The SingleR R/Bioconductor package (v4.6.0) was used to annotate cell type at cell level with the mouse Immunological Genome Project (*37*) obtained through the celldex R/Bioconductor package (v4.6.0).

Differential gene expression analysis between lung and liver tissue and metastatic niches of MVT1 was performed after batch integration and pseudobulk aggregation at mouse level. Differential gene expression was computed using quasi-likelihood F-tests from edgeR R/Bioconductor package (v1.38.3) with robust dispersion estimates. Gene set enrichment analysis (GSEA) was conducted with the fast GSEA method implemented in the R/Bioconductor package clusterProfiler (v4.6.0). As input for GSEA, we used a list of genes ranked by fold-change.

### Cytokine ligand-receptor interaction

We predicted cytokine ligand-receptor interactions between cancer and niche cells using two complementary approaches. First, we followed a similar approach as Kumar et al. (2018) (*38*). We obtained a database of ligand-receptors from SingleCellSignalR (v1.10.0) (*21*) and converted human genes to mouse orthologs with biomaRt (v2.54.0) using Ensembl (v.105). To curate the list of interactions we selected cytokine ligands included in the Gene Ontology term “cytokine activity” (<u>GO:0005125</u>) and filtered out those cytokines and receptors expressed (threshold of 0 counts) in less than 1% of the cells. Finally, for each replicate and organ, we scored interactions using the product of average cytokine normalized expression across all niche cells and the average receptor normalized expression across all cancer cells. We computed two-sided *P* values for the interaction score using permutation test with 1’000 repetitions permuting receptor labels. We selected those interactions with *P* < 0.05 in at least one replicate. To complement the ligand-receptor analysis, we employed CellPhoneDB v3.1.0 (*22*) using database v4.0.0 and the “statistical_analysis” method. The input data contained the normalized expression matrix with mouse genes converted into human orthologs (see details above) and we ran one analysis for each site and replicate. The results table was curated selecting only those interactions involving cytokine ligands in niche cells and their respective receptors in cancer cells.

### Data analysis

Data analysis, statistical testing and visualization were conducted in R (version 4.2.2; R Foundation for Statistical Computing), Bioconductor (v.3.16) and GraphPad Prism 10.

## Data availability

RNA-seq data have been deposited in the Gene Expression Omnibus (GEO, National Center for Biotechnology Information; accession numbers GSE252507, GSE288915 and GSE288916).

## Acknowledgments

We thank members and collaborators of the Aceto Laboratory for scientific feedback and discussions. We thank the Functional Genomic Center Zurich for generating libraries and carrying out next-generation sequencing; the Flow Cytometry Core facility (ETH Zurich) for assistance with FACS-based experiments; A.Offinger (University of Basel) and her team as well as the EPIC team (ETH Zurich) for support with animal work. . The Aceto laboratory is supported by the European Research Council (101001652), the strategic focus area of personalized health and related technologies at ETH Zurich (PHRT-541 and PHRT-960), the Swiss National Science Foundation (212183 and 231353), the Swiss Cancer League (KLS-5636-08-2022), the ETH Lymphoma Challenge (LC-02-22) and ETH Zurich. M.K. Sznurkowska is an EACR-AstraZeneca Fellow, H2020 Marie Skłodowska-Curie Actions (847012) and an EMBO Long Term Fellow (ALTF 899-2018). Y.W.Z. is supported by EMBO Postdoctoral Fellowship (ALTF 142-2023).

## Author contributions

M.K.S. and N.A. designed the study. M.K.S performed all experiments. F.C.G performed all computational analyses and Y.Z performed barcode computational analysis. I.K. performed immunostaining. L.L.O contributed to CTC quantification. I.D. contributed to immunostaining and *in vivo* experiments. A.R., F.D.S, F.A. and K.D.M. collected clinical data. Y.W.Z contributed to seahorse experiments. J.Z. contributed to cytokine treatment experiments. K.S. contributed to *in vivo* experiments. S.B. and M.V. performed tissue sectioning and staining optimisation. M.S. provided MVT1-Thy1.2 cell line. Z.K., S.B. contributed to knockout cell generation. G.P. performed tissue preservation post rapid autopsy. G.A and C.G. assisted with experiments. V.T. supervised some experiments. I.M. provided the construct for niche-labelling. H.F.H., A.W. and M.V. supervised clinical data acquisition. M.L. performed and supervised rapid autopsy. M.K.S, A.G. and N.A. wrote the manuscript. All authors read the manuscript and provided feedback.

## Competing interests

N.A. is a co-founder and member of the board of PAGE Therapeutics AG, Switzerland, listed as an inventor in patent applications related to CTCs, a paid consultant for companies with an interest in liquid biopsies, and a Novartis shareholder. All other authors declare no competing interests.

## Additional information

**Extended Data** Fig. 1 **| Disease outcome data for breast cancer patients diagnosed with initial mono-metastasis in liver, lung, and bone.**

**A,B.** Kaplan-Meier plots comparing time to next event (metastasis or death) (*left*) and overall survival (*right*) of breast cancer patients with mono-metastatic liver (**A**, *n* = 10) and lung (**B**, *n* = 12) lesions *versus* patients with mono-metastatic bone lesions (*n* = 58). **C-E.** Kaplan-Meier plots (*left*) and box plots (*right*) depicting the time to next event (metastasis or death) for SEER cohorts having either liver or lung initial mono-metastasis involvement for TNBC (**C**), colorectal (**D**) and pancreatic ductal adenocarcinoma (PDAC) (**E**) cancers. *n* = 278 (TNBC), *n* = 13’355 (colorectal), *n* = 26’680 (PDAC) for liver metastasis, and *n* = 606 (TNBC), *n* = 1’864 (colorectal), *n* = 2’986 (PDAC) for lungs metastasis. (**C-E**) Each point represents an individual patient, and the cross bars represent the median. *P* values were calculated using the two-sided Wilcoxon rank-sum test.

**Extended Data Fig. 2 | Barcode complexity in primary tumor, lung and liver metastases.**

**A.** Counts of barcodes detected in MVT1 primary tumor (PT), lung and liver metastases (M-lung, M-liver, respectively), as well as CTCs using a threshold of five counts per million (CPM) per sample. The lines indicate the mean along with standard deviation. **B-D.** Dot plots showing relative abundance (percentage) of distinct M-lung-(*left*) and M-liver-unique (*right*) barcodes in individual mice; mouse A (**B**), B (**C**) and C (**D**) are shown, across all metastatic sites.

**Extended Data Fig. 3 | Unique cellular barcodes in lung and liver metastases.**

**A-D**. Volcano plots illustrating the unique barcodes for lung and liver metastases (M-lung, M-liver, respectively) in individual mice carrying an orthotopic MVT1-GFP-luciferase tumor. Mouse B (**A, B**) and mouse C (**C, D**) are shown. **E-G.** Heatmaps displaying unique lung (*top,* M-lung) and liver (*bottom,* M-liver) metastases barcode levels, represented separately for each mouse. The color depicts standardized Z-score. **H-I.** Principle component analysis (PCA) plot of 500 top variable barcodes represented in the primary tumor (PT), circulating tumor cells (CTCs), lung (M-lung) and liver (M-liver) metastases of individual mice.

**Extended Data Fig. 4 | Enriched reactome pathways and Gene Ontology (GO)-biological process terms in the MVT1-GFP-luciferase orthotopic model.**

**A,B**. Enriched reactome pathways and GO-biological processes (adjusted *P* value < 0.01) for MVT1 cells associated with a given organ (**A**) and organ-specific gene expression clusters (**B**). Top 20 pathways for each group with hypergeometric test adjusted *P* value < 0.01 are shown. PT, M-lung, M-liver correspond to primary tumor, lung metastases, and liver metastases, respectively. Displayed is a condensed set of ontologies following simplification through semantic similarity. A complete pathway list is included in Supplementary Tables 3 and 4.

**Extended Data Fig. 5 | Enriched reactome pathways and Gene Ontology (GO)-biological process terms in the BR16 CTC-derived breast cancer orthotopic model.**

**A.** Schematic representing the experimental design of the single-cell RNA-sequencing experiment in NSG mice carrying orthotopic BR16-GFP-luciferase tumors. **B.** Uniform Manifold Approximation and Projection (UMAP) embedding colored by organ of origin, i.e. primary tumor (PT), lung metastases (M-lung) and liver metastases (M-liver). **C.** Enriched reactome pathways and GO-biological process terms in organ specific gene expression clusters. Top 20 pathways for each group with hypergeometric test adjusted *P* < 0.01 are shown. PT, M-lung, M-liver correspond to primary tumor, lung metastases, and liver metastases, respectively. Displayed is a condensed set of ontologies following simplification through semantic similarity. The complete pathway list is provided in Supplementary Table 5.

**Extended Data Fig. 6 | Abundance of non-epithelial cell populations in healthy lung and liver versus lung and liver metastatic niches of tumor-bearing mice.**

Box plot showing the proportion of macrophages, endothelial cells, monocytes, neutrophils, dendritic cells (DC) and fibroblasts in normal tissue and lung (**A**) and liver (**B**) metastatic niches (M-niche). The line indicates median and the box denotes interquartile range. *, *P* < 0.05; **, *P* < 0.01; ***, *P* < 0.001; ****, *P* < 0.0001 by two-sided *t* test.

**Extended Data Fig. 7 | Gene expression profiles of identified niche cell types.**

Dot plot gene expression patterns of macrophages, monocytes, dendritic cells, neutrophils, fibroblasts, and endothelial cells found within the primary tumor (PT-TME), lung and liver metastatic niches (M-niche) are shown. Dot color and size correspond to the gene expression levels of indicated genes and proportion of cells expressing a cell-specific gene, respectively.

**Extended Data Fig. 8 | Gene expression of tissue-specific and general macrophage markers.** Dot plot showing gene expression profile of tissue-specific and general macrophage markers macrophages found within the primary tumor microenvironment (PT-TME), as well as the metastatic niche (M-niche) of lung and liver. Dot color and size correspond to the gene expression levels and proportion of cells expressing a specific marker, respectively.

**Extended Data Fig. 9 | Differential gene expression analysis between healthy lung and liver tissue and metastatic niches in tumor-bearing animals.**

**A-B.** Volcano plot showing genes upregulated (*red;* positive log_2_ fold change) or downregulated (*blue*; negative log_2_ fold change) in the metastatic niche (M-niche) of organs from mice of injected orthotopically with MVT1-GFP-luciferase cells compared to healthy tissue for lung macrophages (**A**) and liver endothelial cells (**B**). **C-D.** Enriched reactome pathways and gene ontology (GO)-biological process terms in macrophages in the lung and liver metastatic niches (M-niche). (**C**) and endothelial cells in the liver metastatic niche (M-niche) (**D**) compared to healthy tissue. Positive normalized enrichment score (NES) corresponds to upregulation in the metastatic niche. Top 20 pathways for each group with hypergeometric test adjusted *P* < 0.01 are shown. Displayed is a condensed set of ontologies following simplification through semantic similarity.

**Extended Data Fig. 10 | Cancer-niche cell interactions in primary tumor, liver, and lung metastases.**

**A.** Dot plot representing interactions (non-normalized) between cytokine ligands (niche cells) and their receptors (cancer cells) in primary tumor (PT), lung and liver metastases. Dot color and size correspond to the interaction scores in individual animals and to the respective *P* value (*P* < 0.05). **B.** Box plot showing the distribution of interaction scores for selected cytokine-receptor pairs in the PT, lung, and liver metastases, with the line indicating the median and the box borders representing interquartile range. *P* value is calculated by two-sided *t* test; *, *P* < 0.05, **, *P* < 0.01. **C**. Dot plot representing selected cytokine-receptor pairs per cell type and anatomical site. Dot color and size correspond to the average interaction score across mouse replicates. DC, dendritic cells.

**Extended Data Fig. 11 | Cytokine gene expression in niche cells and corresponding receptors in cancer cells.**

UMAP displaying cytokine and receptor expression in niche cells (M-niche) and cancer cells, respectively for *Ssp1*-*Itgb1*, *Bmp6*-*Acvr1*/*Bmpr1a*, *Bmpr2* receptor pairs (refer to **Fig.3B** and **Fig. 2B** for the cell type and organ of origin).

**Extended Data Fig. 12 | Cytokine treatment *in vitro*.**

**A.** Western blot analysis of phospho-SMAD1/5 (P-SMAD1/5), total SMAD 1/5 and actin in untreated (control) and BMP2 treated MVT1-GFP-luciferase cells. Treatment time amounted to 15 minutes. **B.** Bar graph showing **P**-SMAD1/5 levels normalized to total SMAD1/5, revealing activation of SMAD1/5 following BMP2 treatment. **, *P* < 0.01 by two-sided *t* test.

**Supplementary Table 1. Clinical characteristics of breast cancer patients within the analyzed cohorts.**

The table shows molecular subtype, age at primary diagnosis, survival status, follow up in days, initial and subsequent metastatic sites. ER, PR, HR stand for estrogen receptor, progesterone receptor and hormone receptors, respectively.

**Supplementary Table 2. Sequence of metastatic diagnoses for breast cancer patients within the analyzed cohorts.**

The table shows detailed information regarding metastasis diagnosis, organ involvement and timing of progression for breast cancer patients included in our study.

**Supplementary Table 3. Complete list of enriched reactome pathways and Gene Ontology – Biological Process (adjusted *P* value < 0.01) for organ specific gene expression clusters associated with the primary tumor (PT), lung and liver metastases in MVT1-GFP-luciferase model.**

The table includes PCA cluster number, pathway ID, pathway description, gene ratio, background ratio (BgRatio), *P*-value, adjusted *P*-value, number of cluster marker genes from a given gene set (count), number of genes in a gene set (set size) and the ratio between *count* and *set size* (gene proportion).

**Supplementary Table 4. Complete list of enriched reactome pathways and Gene Ontology – Biological Process (adjusted *P* value < 0.01) for all cancer cells associated with the primary tumor (PT), lung and liver metastases in MVT1-GFP-luciferase model.**

The table includes PCA cluster number, pathway ID, pathway description, gene ratio, background ratio (BgRatio), *P*-value, adjusted *P*-value, number of cluster marker genes from a given gene set (count), number of genes in a gene set (set size) and the ratio between *count* and *set size* (gene proportion).

**Supplementary Table 5. Complete list of enriched reactome pathways and Gene Ontology – Biological Process (adjusted *P* value < 0.01) for all cancer cells associated with the primary tumor (PT), lung and liver metastases in Br16-GFP-luciferase model.**

The table includes PCA cluster number, pathway ID, pathway description, gene ratio, background ratio (BgRatio), *P*-value, adjusted *P*-value, number of cluster marker genes from a given gene set (count), number of genes in a gene set (set size) and the ratio between *count* and *set size* (gene proportion).

## Notes

### Summary of Updates

Revisions to include new experiments. Fig. 1-4 revised.

## References

1. H. Dillekås, M. S. Rogers, O. Straume, Are 90% of deaths from cancer caused by metastases? Cancer Med. 8, 5574–5576 (2019).

2. P. Gui, T. G. Bivona, Evolution of metastasis: new tools and insights. Trends Cancer 8, 98–109 (2022).

3. A. L. Parker, M. Benguigui, J. Fornetti, E. Goddard, S. Lucotti, J. Insua-Rodríguez, A. P. Wiegmans, Early Career Leadership Council of the Metastasis Research Society, Current challenges in metastasis research and future innovation for clinical translation. Clin. Exp. Metastasis 39, 263–277 (2022).

4. A. A. Abdul Pari, M. Singhal, H. G. Augustin, Emerging paradigms in metastasis research. J. Exp. Med. 218, e20190218 (2021).

5. H. Sung, J. Ferlay, R. L. Siegel, M. Laversanne, I. Soerjomataram, A. Jemal, F. Bray, Global Cancer Statistics 2020: GLOBOCAN Estimates of Incidence and Mortality Worldwide for 36 Cancers in 185 Countries. CA. Cancer J. Clin. 71, 209–249 (2021).

6. H. Kennecke, R. Yerushalmi, R. Woods, M. C. U. Cheang, D. Voduc, C. H. Speers, T. O. Nielsen, K. Gelmon, Metastatic Behavior of Breast Cancer Subtypes. J. Clin. Oncol. 28, 3271–3277 (2010).

7. P. Zagami, L. A. Carey, Triple negative breast cancer: Pitfalls and progress. Npj Breast Cancer 8, 95 (2022).

8. K. H. Allison, M. E. H. Hammond, M. Dowsett, S. E. McKernin, L. A. Carey, P. L. Fitzgibbons, D. F. Hayes, S. R. Lakhani, M. Chavez-MacGregor, J. Perlmutter, C. M. Perou, M. M. Regan, D. L. Rimm, W. F. Symmans, E. E. Torlakovic, L. Varella, G. Viale, T. F. Weisberg, L. M. McShane, A. C. Wolff, Estrogen and Progesterone Receptor Testing in Breast Cancer: American Society of Clinical Oncology/College of American Pathologists Guideline Update. Arch. Pathol. Lab. Med. 144, 545–563 (2020).

9. Q. Wu, J. Li, S. Zhu, J. Wu, C. Chen, Q. Liu, W. Wei, Y. Zhang, S. Sun, Breast cancer subtypes predict the preferential site of distant metastases: a SEER based study. Oncotarget 8, 27990–27996 (2017).

10. H. Zhao, Y. Gong, F. Ye, H. Ling, X. Hu, Incidence and prognostic factors of patients with synchronous liver metastases upon initial diagnosis of breast cancer: a population-based study. Cancer Manag. Res. **Volume** 10, 5937–5950 (2018).

11. L. Gerratana, V. Fanotto, M. Bonotto, S. Bolzonello, A. M. Minisini, G. Fasola, F. Puglisi, Pattern of metastasis and outcome in patients with breast cancer. Clin. Exp. Metastasis 32, 125–133 (2015).

12. W. Xiao, S. Zheng, P. Liu, Y. Zou, X. Xie, P. Yu, H. Tang, X. Xie, Risk factors and survival outcomes in patients with breast cancer and lung metastasis: a population-based study. Cancer Med. 7, 922–930 (2018).

13. H. Pan, R. Gray, J. Braybrooke, C. Davies, C. Taylor, P. McGale, R. Peto, K. I. Pritchard, J. Bergh, M. Dowsett, D. F. Hayes, 20-Year Risks of Breast-Cancer Recurrence after Stopping Endocrine Therapy at 5 Years. N. Engl. J. Med. 377, 1836–1846 (2017).

14. S. J. Lee, S. Park, H. K. Ahn, J. H. Yi, E. Y. Cho, J. M. Sun, J. E. Lee, S. J. Nam, J.-H. Yang, Y. H. Park, J. S. Ahn, Y.-H. Im, Implications of Bone-Only Metastases in Breast Cancer: Favorable Preference with Excellent Outcomes of Hormone Receptor Positive Breast Cancer. Cancer Res. Treat. 43, 89–95 (2011).

15. R. Wang, Y. Zhu, X. Liu, X. Liao, J. He, L. Niu, The Clinicopathological features and survival outcomes of patients with different metastatic sites in stage IV breast cancer. BMC Cancer 19, 1091 (2019).

16. J. L. Warren, A. Mariotto, D. Melbert, D. Schrag, P. Doria-Rose, D. Penson, K. R. Yabroff, Sensitivity of Medicare Claims to Identify Cancer Recurrence in Elderly Colorectal and Breast Cancer Patients. Med. Care 54, e47–e54 (2016).

17. Surveillance, Epidemiology, and End Results (SEER) Program (www.seer.cancer.gov) SEER*Stat Database: Incidence - SEER Research Data, 8 Registries, Nov 2023 Sub (1975-2021) - Linked To County Attributes - Time Dependent (1990-2022) Income/Rurality, 1969-2022 Counties, National Cancer Institute, DCCPS, Surveillance Research Program, released April 2024, based on the November 2023 submission.

18. C. A. Grzelak, E. T. Goddard, E. E. Lederer, K. Rajaram, J. Dai, R. E. Shor, A. R. Lim, J. Kim, S. Beronja, A. P. W. Funnell, C. M. Ghajar, Elimination of fluorescent protein immunogenicity permits modeling of metastasis in immune-competent settings. Cancer Cell 40, 1–2 (2022).

19. L. Ombrato, E. Nolan, I. Kurelac, A. Mavousian, V. L. Bridgeman, I. Heinze, P. Chakravarty, S. Horswell, E. Gonzalez-Gualda, G. Matacchione, A. Weston, J. Kirkpatrick, E. Husain, V. Speirs, L. Collinson, A. Ori, J.-H. Lee, I. Malanchi, Metastatic-niche labelling reveals parenchymal cells with stem features. Nature 572, 603–608 (2019).

20. M. C. Scheidmann, F. Castro-Giner, K. Strittmatter, I. Krol, A. Paasinen-Sohns, R. Scherrer, C. Donato, S. Gkountela, B. M. Szczerba, Z. Diamantopoulou, S. Muenst, T. Vlajnic, L. Kunz, M. Vetter, C. Rochlitz, V. Taylor, C. Giachino, T. Schroeder, R. J. Platt, N. Aceto, An *In Vivo* CRISPR Screen Identifies Stepwise Genetic Dependencies of Metastatic Progression. Cancer Res. 82, 681–694 (2022).

21. S. Cabello-Aguilar, M. Alame, F. Kon-Sun-Tack, C. Fau, M. Lacroix, J. Colinge, SingleCellSignalR: inference of intercellular networks from single-cell transcriptomics. Nucleic Acids Res. 48, e55–e55 (2020).

22. L. Garcia-Alonso, L.-F. Handfield, K. Roberts, K. Nikolakopoulou, R. C. Fernando, L. Gardner, B. Woodhams, A. Arutyunyan, K. Polanski, R. Hoo, C. Sancho-Serra, T. Li, K. Kwakwa, E. Tuck, V. Lorenzi, H. Massalha, M. Prete, V. Kleshchevnikov, A. Tarkowska, T. Porter, C. I. Mazzeo, S. Van Dongen, M. Dabrowska, V. Vaskivskyi, K. T. Mahbubani, J. Park, M. Jimenez-Linan, L. Campos, V. Yu. Kiselev, C. Lindskog, P. Ayuk, E. Prigmore, M. R. Stratton, K. Saeb-Parsy, A. Moffett, L. Moore, O. A. Bayraktar, S. A. Teichmann, M. Y. Turco, R. Vento-Tormo, Mapping the temporal and spatial dynamics of the human endometrium in vivo and in vitro. Nat. Genet. 53, 1698– 1711 (2021).

23. K. Lavery, P. Swain, D. Falb, M. H. Alaoui-Ismaili, BMP-2/4 and BMP-6/7 Differentially Utilize Cell Surface Receptors to Induce Osteoblastic Differentiation of Human Bone Marrow-derived Mesenchymal Stem Cells. J. Biol. Chem. 283, 20948– 20958 (2008).

24. M. Omi, V. Kaartinen, Y. Mishina, Activin A receptor type 1–mediated BMP signaling regulates RANKL-induced osteoclastogenesis via canonical SMAD-signaling pathway. J. Biol. Chem. 294, 17818–17836 (2019).

25. C. C. Ho, D. J. Bernard, Bone Morphogenetic Protein 2 Signals via BMPR1A to Regulate Murine Follicle-Stimulating Hormone Beta Subunit Transcription1. Biol. Reprod. 81, 133–141 (2009).

26. L. Bai, H.-M. Chang, J.-C. Cheng, C. Klausen, G. Chu, P. C. K. Leung, G. Yang, SMAD1/5 mediates bone morphogenetic protein 2-induced up-regulation of BAMBI expression in human granulosa-lutein cells. Cell. Signal. 37, 52–61 (2017).

27. W. Zhang, I. L. Bado, J. Hu, Y.-W. Wan, L. Wu, H. Wang, Y. Gao, H.-H. Jeong, Z. Xu, X. Hao, B. M. Lege, R. Al-Ouran, L. Li, J. Li, L. Yu, S. Singh, H. C. Lo, M. Niu, J. Liu, W. Jiang, Y. Li, S. T. C. Wong, C. Cheng, Z. Liu, X. H.-F. Zhang, The bone microenvironment invigorates metastatic seeds for further dissemination. Cell 184, 2471–2486.e20 (2021).

28. Y. Yang, H. H. Yang, Y. Hu, P. H. Watson, H. Liu, T. R. Geiger, M. R. Anver, D. C. Haines, P. Martin, J. E. Green, M. P. Lee, K. W. Hunter, L. M. Wakefield, Immunocompetent mouse allograft models for development of therapies to target breast cancer metastasis. Oncotarget 8, 30621–30643 (2017).

29. M. Yu, A. Bardia, N. Aceto, F. Bersani, M. W. Madden, M. C. Donaldson, R. Desai, H. Zhu, V. Comaills, Z. Zheng, B. S. Wittner, P. Stojanov, E. Brachtel, D. Sgroi, R. Kapur, T. Shioda, D. T. Ting, S. Ramaswamy, G. Getz, A. J. Iafrate, C. Benes, M. Toner, S. Maheswaran, D. A. Haber, Ex vivo culture of circulating breast tumor cells for individualized testing of drug susceptibility. Science 345, 216–220 (2014).

30. M. Martin, Cutadapt removes adapter sequences from high-throughput sequencing reads. EMBnet.journal 17, 10–12 (2011).

31. B. Langmead, S. L. Salzberg, Fast gapped-read alignment with Bowtie 2. Nat. Methods 9, 357–359 (2012).

32. P. Danecek, J. K. Bonfield, J. Liddle, J. Marshall, V. Ohan, M. O. Pollard, A. Whitwham, T. Keane, S. A. McCarthy, R. M. Davies, H. Li, Twelve years of SAMtools and BCFtools. GigaScience 10, giab008 (2021).

33. M. E. Ritchie, B. Phipson, D. Wu, Y. Hu, C. W. Law, W. Shi, G. K. Smyth, limma powers differential expression analyses for RNA-sequencing and microarray studies. Nucleic Acids Res. 43, e47–e47 (2015).

34. H.-J. Zhao, H.-M. Chang, C. Klausen, H. Zhu, Y. Li, P. C. K. Leung, Bone morphogenetic protein 2 induces the activation of WNT/β-catenin signaling and human trophoblast invasion through up-regulating BAMBI. Cell. Signal. 67, 109489 (2020).

35. J. Chen, H. Liu, J. Liu, J. Qi, B. Wei, J. Yang, H. Liang, Y. Chen, J. Chen, Y. Wu, L. Guo, J. Zhu, X. Zhao, T. Peng, Y. Zhang, S. Chen, X. Li, D. Li, T. Wang, D. Pei, H3K9 methylation is a barrier during somatic cell reprogramming into iPSCs. Nat. Genet. 45, 34–42 (2013).

36. B. Phipson, C. B. Sim, E. R. Porrello, A. W. Hewitt, J. Powell, A. Oshlack, propeller: testing for differences in cell type proportions in single cell data.

37. The Immunological Genome Project Consortium, T. S. P. Heng, M. W. Painter, K. Elpek, V. Lukacs-Kornek, N. Mauermann, S. J. Turley, D. Koller, F. S. Kim, A. J. Wagers, N. Asinovski, S. Davis, M. Fassett, M. Feuerer, D. H. D. Gray, S. Haxhinasto, J. A. Hill, G. Hyatt, C. Laplace, K. Leatherbee, D. Mathis, C. Benoist, R. Jianu, D. H. Laidlaw, J. A. Best, J. Knell, A. W. Goldrath, J. Jarjoura, J. C. Sun, Y. Zhu, L. L. Lanier, A. Ergun, Z. Li, J. J. Collins, S. A. Shinton, R. R. Hardy, R. Friedline, K. Sylvia, J. Kang, The Immunological Genome Project: networks of gene expression in immune cells. Nat. Immunol. 9, 1091–1094 (2008).

38. M. P. Kumar, J. Du, G. Lagoudas, Y. Jiao, A. Sawyer, D. C. Drummond, D. A. Lauffenburger, A. Raue, Analysis of Single-Cell RNA-Seq Identifies Cell-Cell Communication Associated with Tumor Characteristics. Cell Rep. 25, 1458–1468.e4 (2018).

