## Supplementary Figures for "Charting the liver and lung metastatic niche in breast cancer"

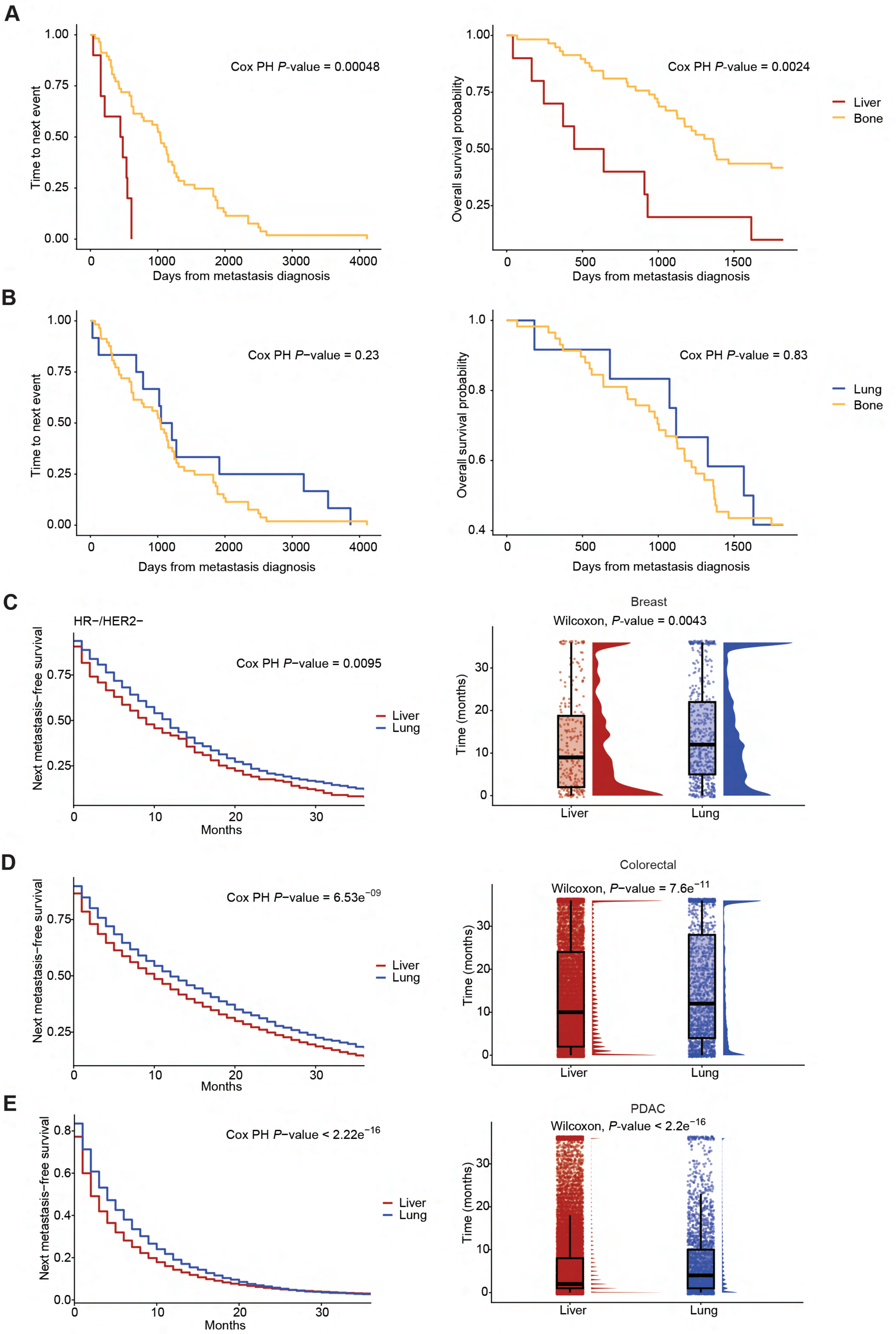


**Extended Data Fig. 1 | Disease outcome data for breast cancer patients diagnosed with initial mono-metastasis in liver, lung, and bone.** **A,B.** Kaplan-Meier plots comparing time to next event (metastasis or death) (*left*) and overall survival (*right*) of breast cancer patients with mono-metastatic liver (**A**, $n = 10$) and lung (**B**, $n = 12$) lesions *versus* patients with mono-metastatic bone lesions ($n = 58$). **C-E.** Kaplan-Meier plots (*left*) and box plots (*right*) depicting the time to next event (metastasis or death) for SEER cohorts having either liver or lung initial mono-metastasis involvement for TNBC (**C**), colorectal (**D**) and pancreatic ductal adenocarcinoma (PDAC) (**E**) cancers. $n = 278$ (TNBC), $n = 13355$ (colorectal), $n = 26680$ (PDAC) for liver metastasis, and $n = 606$ (TNBC), $n = 1864$ (colorectal), $n = 2986$ (PDAC) for lungs metastasis. (**C-E**) Each point represents an individual patient, and the cross bars represent the median. *P* values were calculated using the one-sided Wilcoxon rank-sum test.

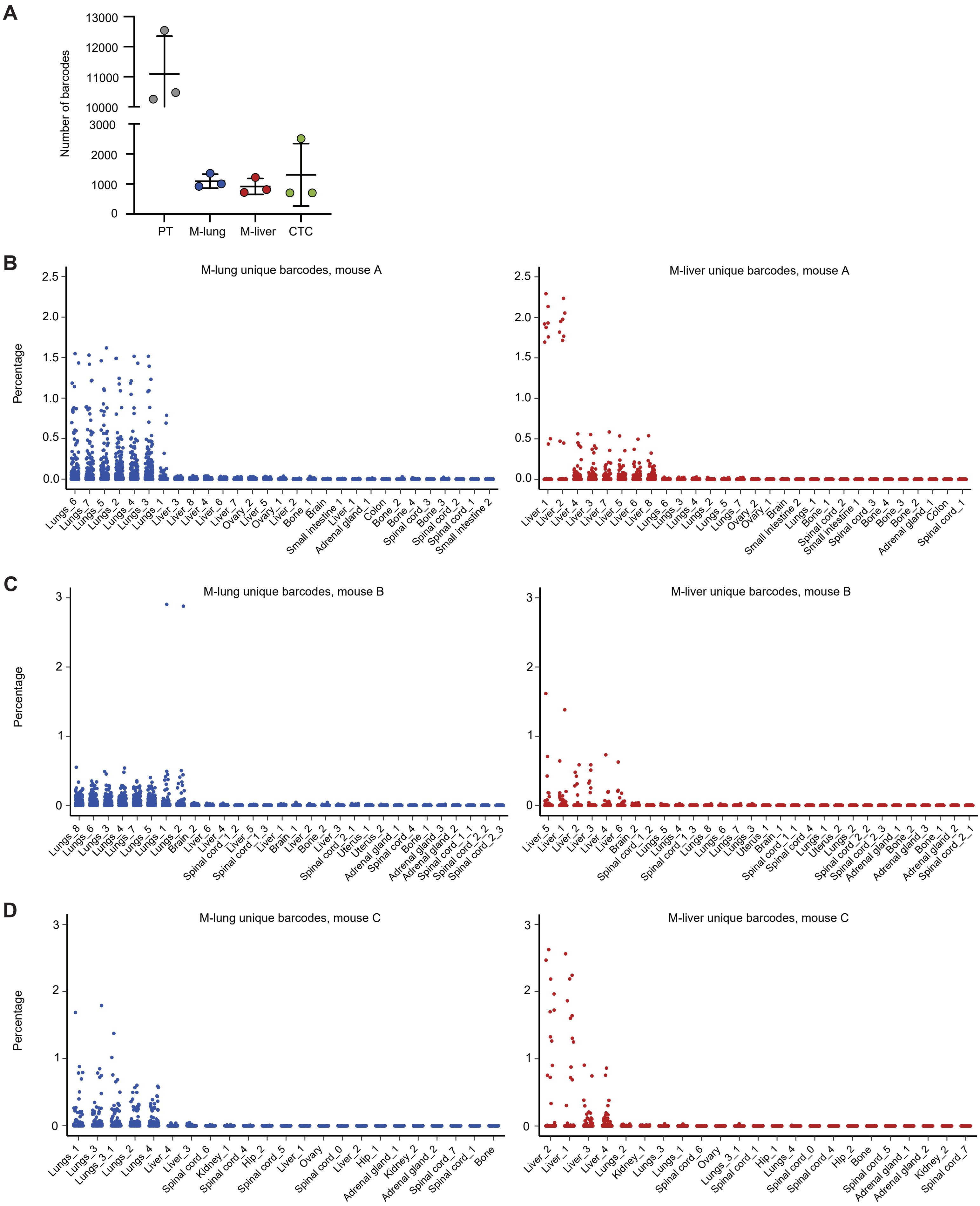

**Extended Data Fig. 2 | Barcode complexity in primary tumor, lung and liver metastases.**
**A.** Counts of barcodes detected in MVT1 primary tumor (PT), lung and liver metastases (M-lung, M-liver, respectively), as well as CTCs using a threshold of five counts per million (CPM) per sample. The lines indicate the mean along with standard deviation. **B-D.** Dot plots showing relative abundance (percentage) of distinct M-lung- (*left*) and M-liver-unique (*right*) barcodes in individual mice; mouse A (**B**), B (**C**) and C (**D**) are shown, across all metastatic sites.

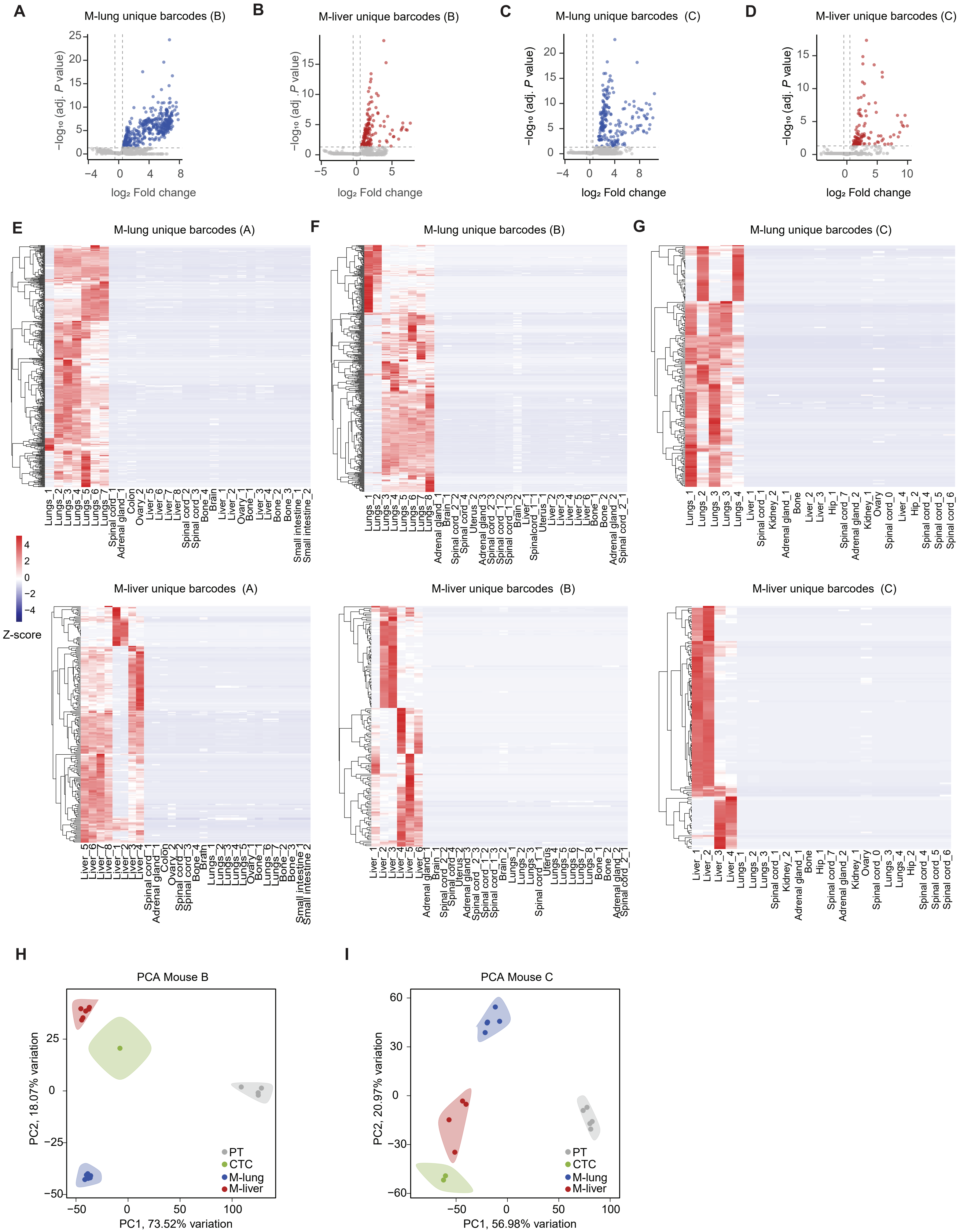


**Extended Data Fig. 3 | Unique cellular barcodes in lung and liver metastases.**
**A-D**. Volcano plots illustrating the unique barcodes for lung and liver metastases (M-lung, M-liver, respectively) in individual mice carrying an orthotopic MVT1-GFP-luciferase tumor. Mouse B (**A, B**) and mouse C (**C, D**) are shown. **E-G.** Heatmaps displaying unique lung (*top,* M-lung) and liver (*bottom*, M-liver) metastases barcode levels, represented separately for each mouse. The color depicts standardized Z-score. **H-I.** Principle component analysis (PCA) plot of 500 top variable barcodes represented in the primary tumor (PT), circulating tumor cells (CTCs), lung (M-lung) and liver (M-liver) metastases of individual mice.

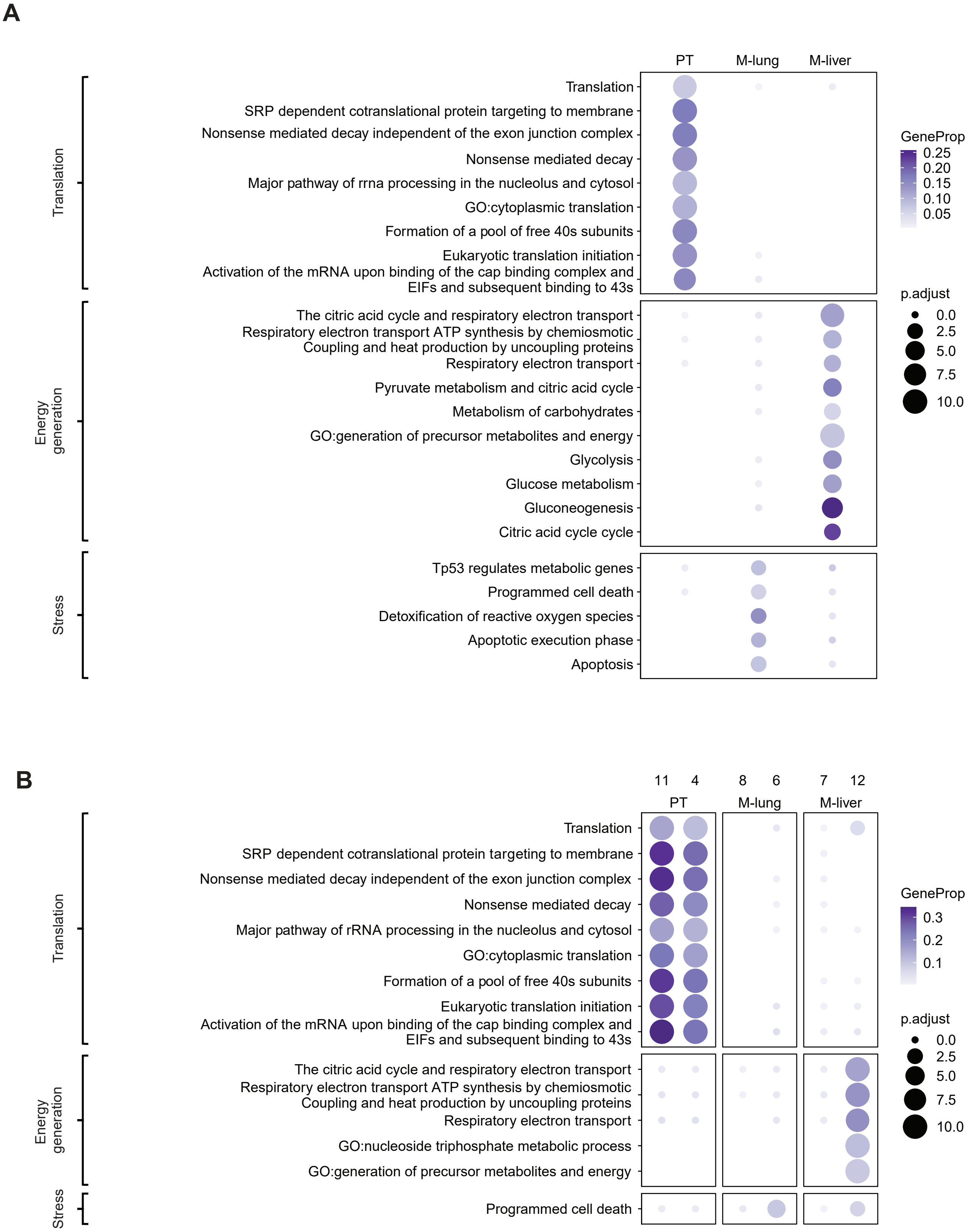


**Extended Data Fig. 4 | Enriched reactome pathways and Gene Ontology (GO)-biological process terms in the MVT1-GFP-luciferase orthotopic model.** **A,B**. Enriched reactome pathways and GO-biological processes (adjusted *P* value < 0.01) for MVT1 cells associated with a given organ (**A**) and organ-specific gene expression clusters (**B**). Top 20 pathways for each group with hypergeometric test adjusted *P* value < 0.01 are shown. PT, M-lung, M-liver correspond to primary tumor, lung metastases, and liver metastases, respectively. Displayed is a condensed set of ontologies following simplification through semantic similarity. A complete pathway list is included in Supplemntary Tables 3 and 4.

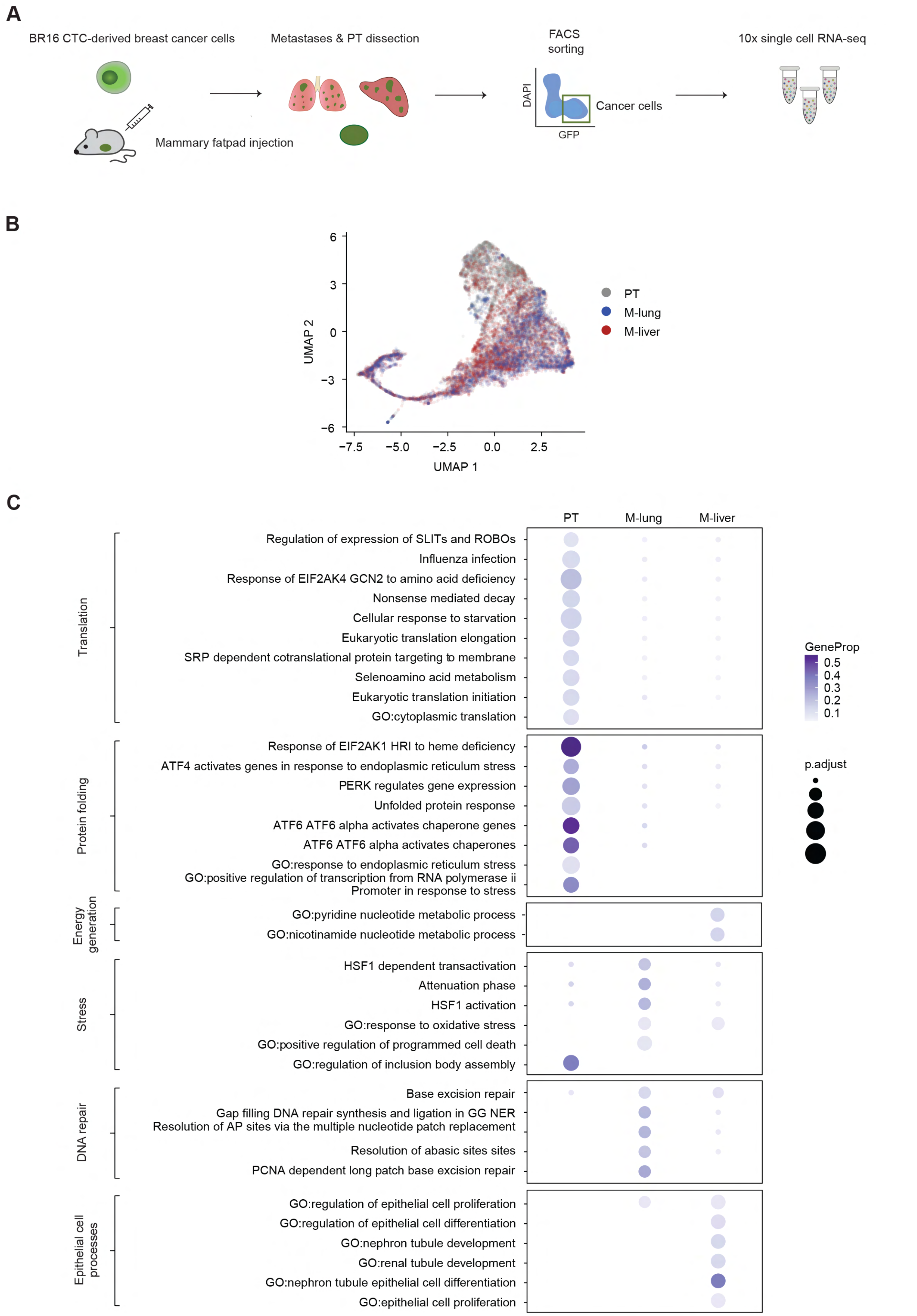


**Extended Data Fig. 5 | Enriched reactome pathways and Gene Ontology (GO)-biological process terms in the BR16 CTC-derived breast cancer orthotopic model.** **A.** Schematic representing the experimental design of the single-cell RNA-sequencing experiment in NSG mice carrying orthotopic BR16-GFP-luciferase tumors. **B.** Uniform Manifold Approximation and Projection (UMAP) embedding colored by organ of origin, i.e. primary tumor (PT), lung metastases (M-lung) and liver metastases (M-liver). **C.** Enriched reactome pathways and GO-biological process terms in organ specific gene expression clusters. Top 20 pathways for each group with hypergeometric test adjusted $P$ < 0.01 are shown. PT, M-lung, M-liver correspond to primary tumor, lung metastases, and liver metastases, respectively. Displayed is a condensed set of ontologies following simplification through semantic similarity. The complete pathway list is provided in Supplementary Table 5.

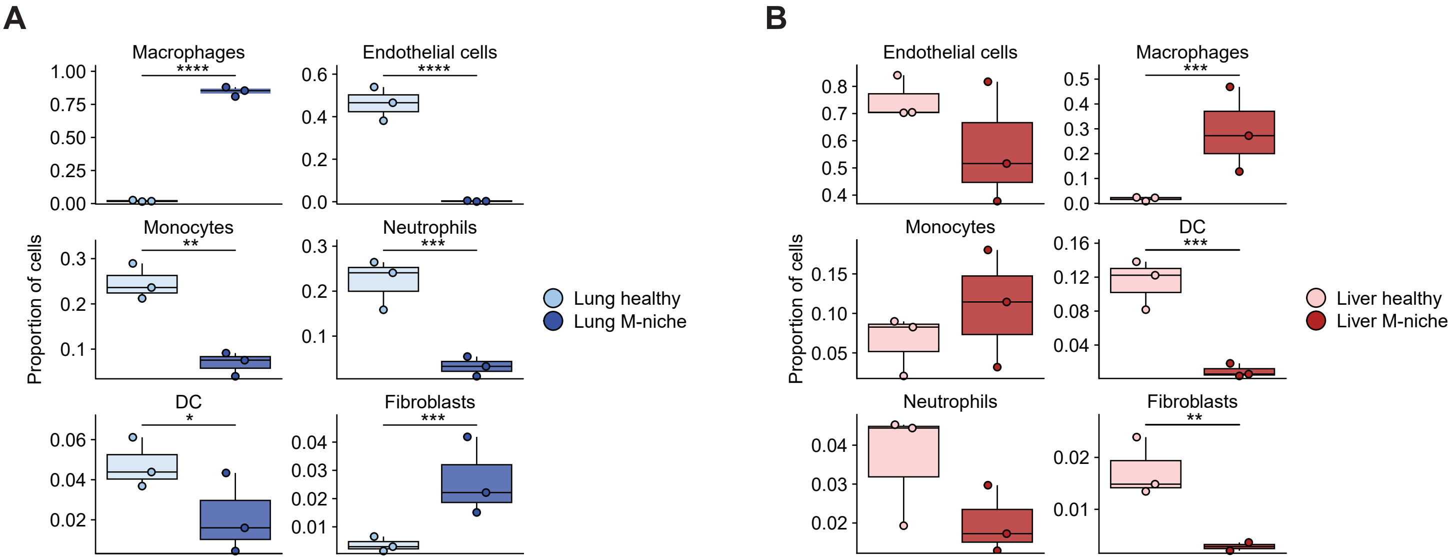


**Extended Data Fig. 6 | Abundance of non-epithelial cell populations in healthy lung and liver versus lung and liver metastatic niches of tumor-bearing mice.** Box plot showing the proportion of macrophages, endothelial cells, monocytes, neutrophils, dendritic cells (DC) and fibroblasts in normal tissue and lung (**A**) and liver (**B**) metastatic niches (M-niche). The line indicates median and the box denotes standard deviation. *, $P < 0.05$; **, $P < 0.01$; ***, $P < 0.001$; ****, $P < 0.0001$ by two-sided *t* test.

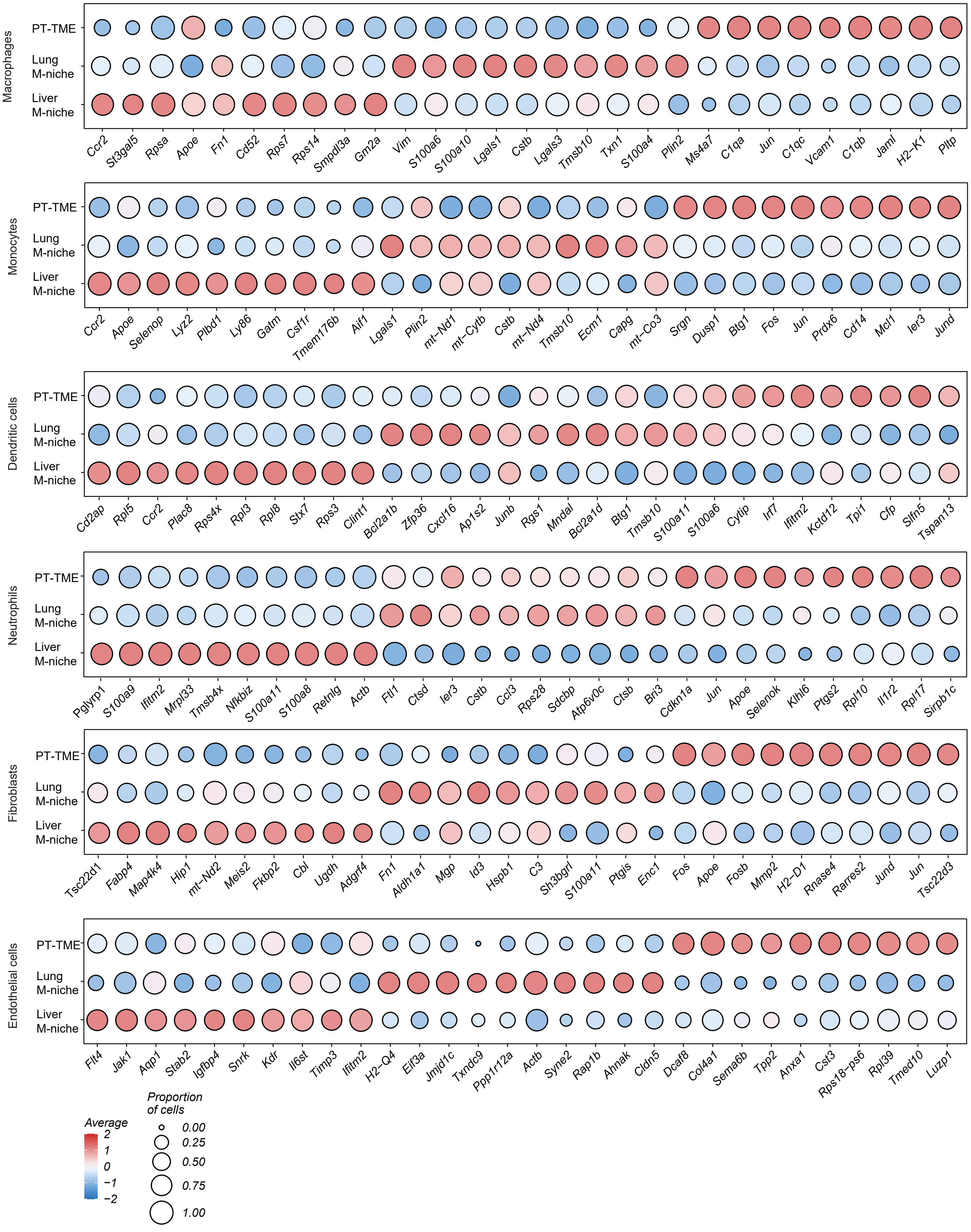


**Extended Data Fig. 7 | Gene expression profiles of identified niche cell types.** Dot plot gene expression patterns of macrophages, monocytes, dendritic cells, neutrophils, fibroblasts, and endothelial cells found within the primary tumor (PT-TME), lung and liver metastatic niches (M-niche) are shown. Dot color and size correspond to the gene expression levels of indicated genes and proportion of cells expressing a cell-specific gene, respectively.

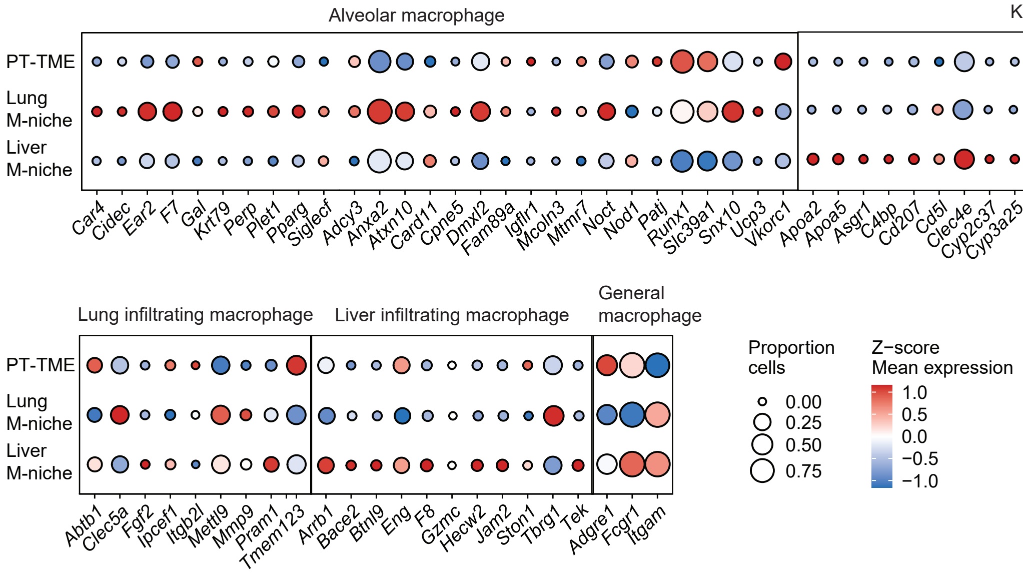


**Extended Data Fig. 8 | Gene expression of tissue-specific and general macrophage markers.** Dot plot showing gene expression profile of tissue-specific and general macrophage markers macrophages found within the primary tumor microenvironment (PT-TME), as well as the metastatic niche (M-niche) of lung and liver. Dot color and size correspond to the gene expression levels and proportion of cells expressing a specific marker, respectively.

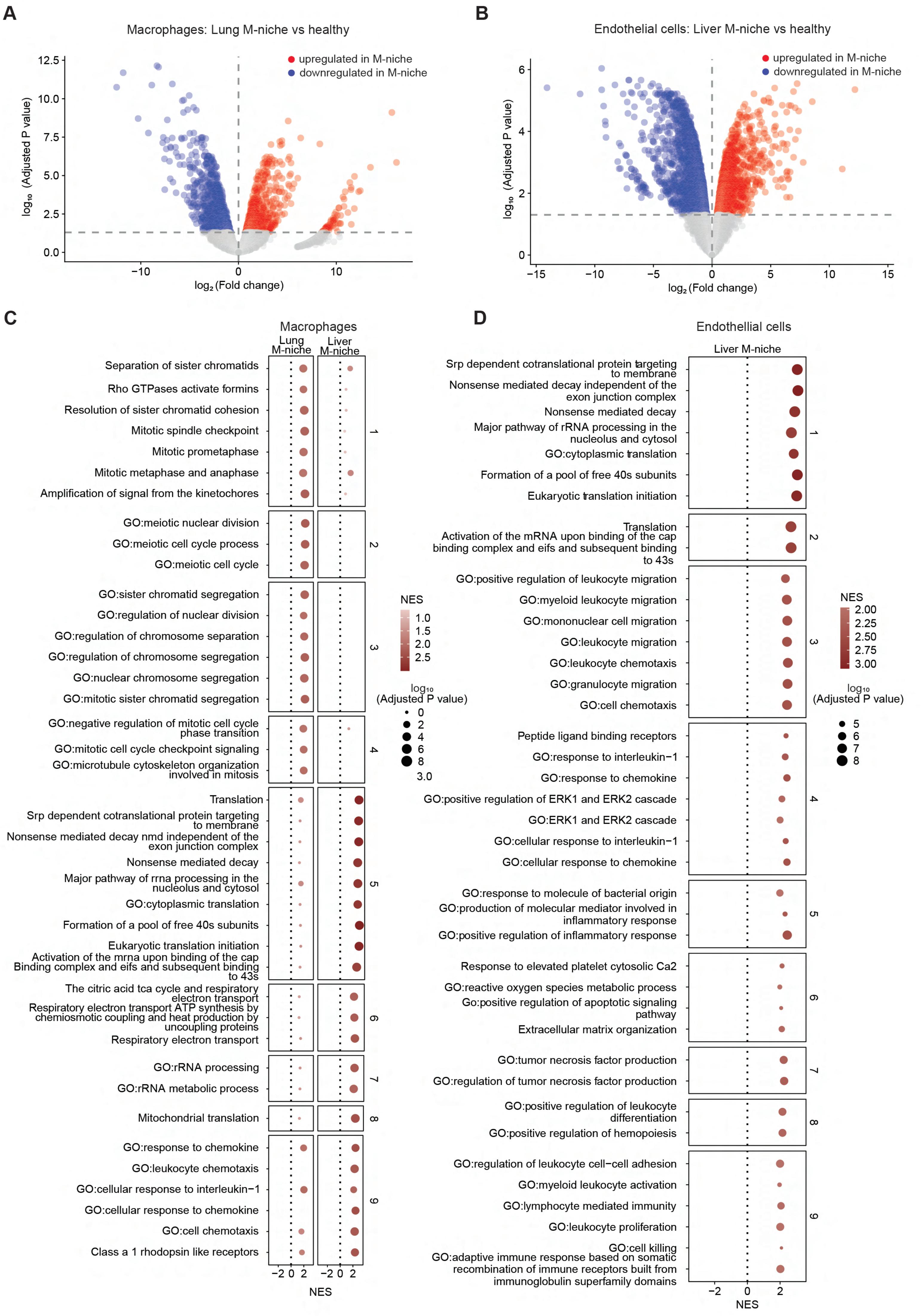

A
Macrophages: Lung M-niche vs healthy
upregulated in M-niche
downregulated in M-niche
log10 (Adjusted P value)
log2 (Fold change)
B
Endothelial cells: Liver M-niche vs healthy
upregulated in M-niche
downregulated in M-niche
log10 (Adjusted P value)
log2 (Fold change)
C
Macrophages
Lung M-niche
Liver M-niche
Separation of sister chromatids
Rho GTPases activate formins
Resolution of sister chromatid cohesion
Mitotic spindle checkpoint
Mitotic prometaphase
Mitotic metaphase and anaphase
Amplification of signal from the kinetochores
GO:meiotic nuclear division
GO:meiotic cell cycle process
GO:meiotic cell cycle
GO:sister chromatid segregation
GO:regulation of nuclear division
GO:regulation of chromosome separation
GO:regulation of chromosome segregation
GO:nuclear chromosome segregation
GO:mitotic sister chromatid segregation
GO:negative regulation of mitotic cell cycle phase transition
GO:mitotic cell cycle checkpoint signaling
GO:microtubule cytoskeleton organization involved in mitosis
Translation
Srp dependent cotranslational protein targeting to membrane
Nonsense mediated decay nmd independent of the exon junction complex
Nonsense mediated decay
Major pathway of rrna processing in the nucleolus and cytosol
GO:cytoplasmic translation
Formation of a pool of free 40s subunits
Eukaryotic translation initiation
Activation of the mrna upon binding of the cap Binding complex and eifs and subsequent binding to 43s
The citric acid tca cycle and respiratory electron transport
Respiratory electron transport ATP synthesis by chemiosmotic coupling and heat production by uncoupling proteins
Respiratory electron transport
GO:rRNA processing
GO:rRNA metabolic process
Mitochondrial translation
GO:response to chemokine
GO:leukocyte chemotaxis
GO:cellular response to interleukin−1
GO:cellular response to chemokine
GO:cell chemotaxis
Class a 1 rhodopsin like receptors
NES
log10 (Adjusted P value)
D
Endothellial cells
Liver M-niche
Srp dependent cotranslational protein targeting to membrane
Nonsense mediated decay independent of the exon junction complex
Nonsense mediated decay
Major pathway of rRNA processing in the nucleolus and cytosol
GO:cytoplasmic translation
Formation of a pool of free 40s subunits
Eukaryotic translation initiation
Translation
Activation of the mRNA upon binding of the cap binding complex and eifs and subsequent binding to 43s
GO:positive regulation of leukocyte migration
GO:myeloid leukocyte migration
GO:mononuclear cell migration
GO:leukocyte migration
GO:leukocyte chemotaxis
GO:granulocyte migration
GO:cell chemotaxis
Peptide ligand binding receptors
GO:response to interleukin−1
GO:response to chemokine
GO:positive regulation of ERK1 and ERK2 cascade
GO:ERK1 and ERK2 cascade
GO:cellular response to interleukin−1
GO:cellular response to chemokine
GO:response to molecule of bacterial origin
GO:production of molecular mediator involved in inflammatory response
GO:positive regulation of inflammatory response
Response to elevated platelet cytosolic Ca2
GO:reactive oxygen species metabolic process
Go:positive regulation of apoptotic signaling pathway
Extracellular matrix organization
GO:tumor necrosis factor production
GO:regulation of tumor necrosis factor production
GO:positive regulation of leukocyte differentiation
GO:positive regulation of hemopoiesis
GO:regulation of leukocyte cell−cell adhesion
GO:myeloid leukocyte activation
GO:lymphocyte mediated immunity
GO:leukocyte proliferation
GO:cell killing
GO:adaptive immune response based on somatic recombination of immune receptors built from immunoglobulin superfamily domains
NES
log10 (Adjusted P value)

**Extended Data Fig. 9 | Differential gene expression analysis between healthy lung and liver tissue and metastatic niches in tumor-bearing animals.**
**A-B.** Volcano plot showing genes upregulated (*red;* positive $log_2$ fold change) or downregulated (*blue*; negative $log_2$ fold change) in the metastatic niche (M-niche) of organs from mice of injected orthotopically with MVT1-GFP-luciferase cells compared to healthy tissue for lung macrophages (**A**) and liver endothelial cells (**B**). **C-D.** Enriched reactome pathways and gene ontology (GO)-biological process terms in macrophages in the lung and liver metastatic niches (M-niche). (**C**) and endothelial cells in the liver metastatic niche (M-niche) (**D**) compared to healthy tissue. Positive normalized enrichment score (NES) corresponds to upregulation in the metastatic niche. Top 20 pathways for each group with hypergeometric test adjusted $P < 0.01$ are shown. Displayed is a condensed set of ontologies following simplification through semantic similarity.

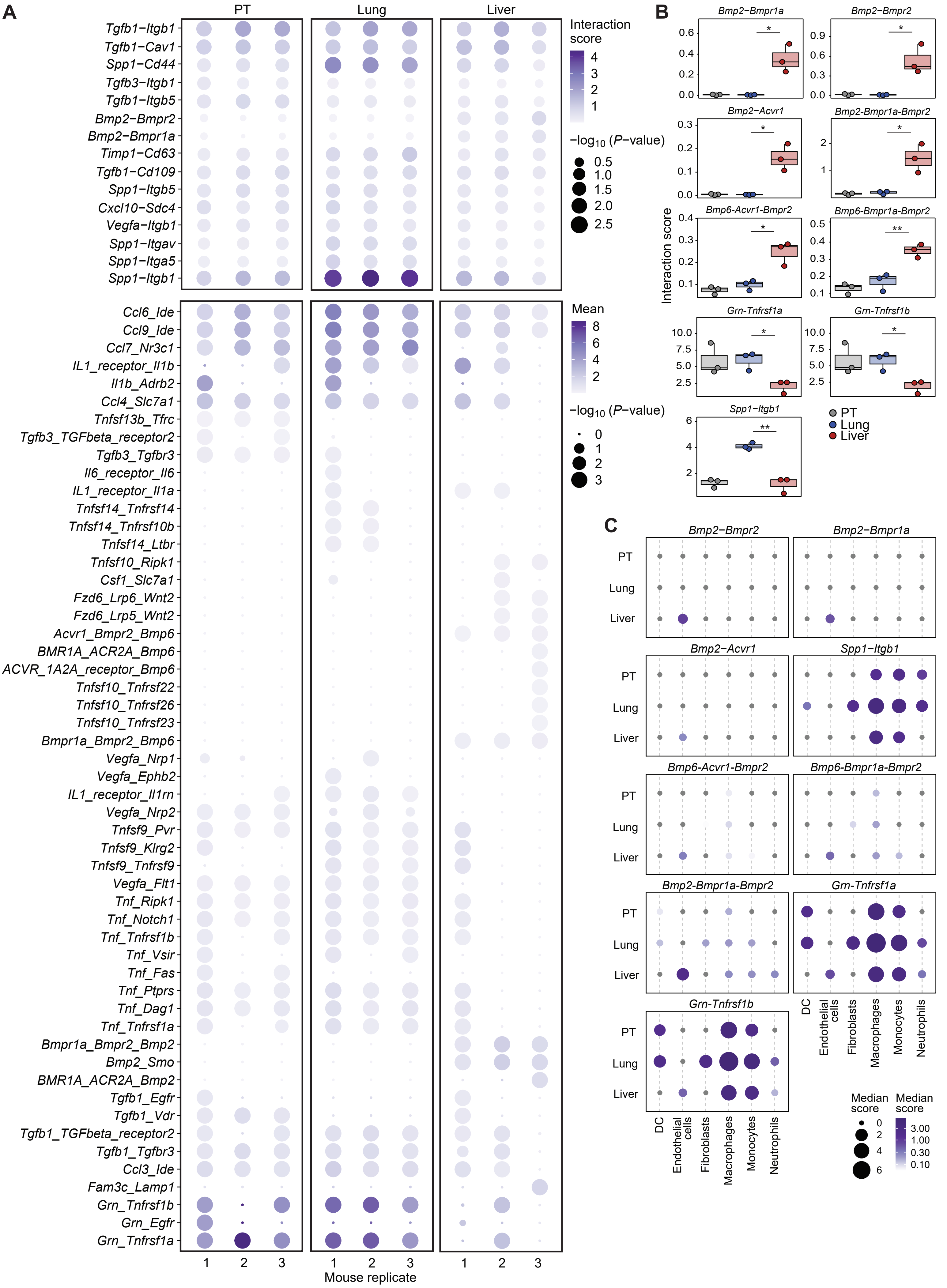
A
PT
Lung
Liver
Tgfb1−Itgb1
Tgfb1−Cav1
Spp1−Cd44
Tgfb3−Itgb1
Tgfb1−Itgb5
Bmp2−Bmpr2
Bmp2−Bmpr1a
Timp1−Cd63
Tgfb1−Cd109
Spp1−Itgb5
Cxcl10−Sdc4
Vegfa−Itgb1
Spp1−Itgav
Spp1−Itga5
Spp1−Itgb1
Interaction score
4
3
2
1
−log10 (P−value)
0.5
1.0
1.5
2.0
2.5
Ccl6_Ide
Ccl9_Ide
Ccl7_Nr3c1
IL1_receptor_Il1b
Il1b_Adrb2
Ccl4_Slc7a1
Tnfsf13b_Tfrc
Tgfb3_TGFbeta_receptor2
Tgfb3_Tgfbr3
Il6_receptor_Il6
IL1_receptor_Il1a
Tnfsf14_Tnfrsf14
Tnfsf14_Tnfrsf10b
Tnfsf14_Ltbr
Tnfsf10_Ripk1
Csf1_Slc7a1
Fzd6_Lrp6_Wnt2
Fzd6_Lrp5_Wnt2
Acvr1_Bmpr2_Bmp6
BMR1A_ACR2A_Bmp6
ACVR_1A2A_receptor_Bmp6
Tnfsf10_Tnfrsf22
Tnfsf10_Tnfrsf26
Tnfsf10_Tnfrsf23
Bmpr1a_Bmpr2_Bmp6
Vegfa_Nrp1
Vegfa_Ephb2
IL1_receptor_Il1rn
Vegfa_Nrp2
Tnfsf9_Pvr
Tnfsf9_Klrg2
Tnfsf9_Tnfrsf9
Vegfa_Flt1
Tnf_Ripk1
Tnf_Notch1
Tnf_Tnfrsf1b
Tnf_Vsir
Tnf_Fas
Tnf_Ptprs
Tnf_Dag1
Tnf_Tnfrsf1a
Bmpr1a_Bmpr2_Bmp2
Bmp2_Smo
BMR1A_ACR2A_Bmp2
Tgfb1_Egfr
Tgfb1_Vdr
Tgfb1_TGFbeta_receptor2
Tgfb1_Tgfbr3
Ccl3_Ide
Fam3c_Lamp1
Grn_Tnfrsf1b
Grn_Egfr
Grn_Tnfrsf1a
Mean
8
6
4
2
−log10 (P−value)
0
1
2
3
1 2 3 1 2 3 1 2 3
Mouse replicate
B
Bmp2−Bmpr1a
Bmp2−Bmpr2
Bmp2−Acvr1
Bmp2-Bmpr1a-Bmpr2
Bmp6-Acvr1-Bmpr2
Bmp6-Bmpr1a-Bmpr2
Grn-Tnfrsf1a
Grn-Tnfrsf1b
Spp1-Itgb1
Interaction score
PT
Lung
Liver
C
Bmp2−Bmpr2
Bmp2−Bmpr1a
Bmp2−Acvr1
Spp1−Itgb1
Bmp6-Acvr1-Bmpr2
Bmp6-Bmpr1a-Bmpr2
Bmp2-Bmpr1a-Bmpr2
Grn-Tnfrsf1a
Grn-Tnfrsf1b
PT
Lung
Liver
DC
Endothelial cells
Fibroblasts
Macrophages
Monocytes
Neutrophils
Median score
0
2
4
6
Median score
3.00
1.00
0.30
0.10

**Extended Data Fig. 10 | Cancer-niche cell interactions in primary tumor, liver, and lung metastases.**
**A.** Dot plot representing interactions (non-normalized) between cytokine ligands (niche cells) and their receptors (cancer cells) in primary tumor (PT), lung and liver metastases. Dot color and size correspond to the interaction scores in individual animals and to the respective $P$ value ($P$ < 0.05). **B.** Box plot showing the distribution of interaction scores for selected cytokine-receptor pairs in the PT, lung, and liver metastases, with the line indicating the median and box borders representing standard deviation. $P$ value is calculated by two-sided $t$ test; *, $P$ < 0.05, **, $P$ < 0.01. **C**. Dot plot representing selected cytokine-receptor pairs per cell type and anatomical site. Dot color and size correspond to the average interaction score across mouse replicates. DC, dendritic cells.

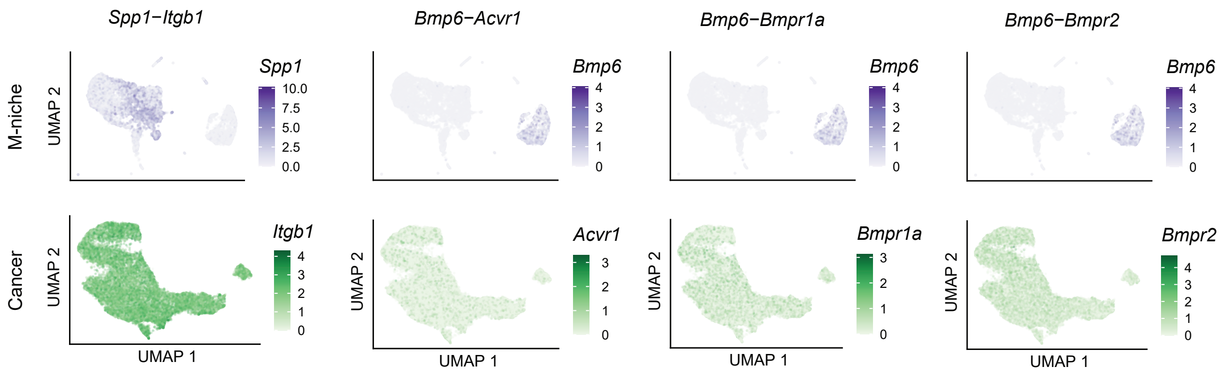


**Extended Data Fig. 11 | Cytokine gene expression in niche cells and corresponding receptors in cancer cells.**
UMAP displaying cytokine and receptor expression in niche cells (M-niche) and cancer cells, respectively for *Ssp1-Itgb1*, *Bmp6-Acvr1*/*Bmpr1a*, *Bmpr2* receptor pairs (refer to **Fig.3B** and **Fig. 2B** for the cell type and organ of origin).

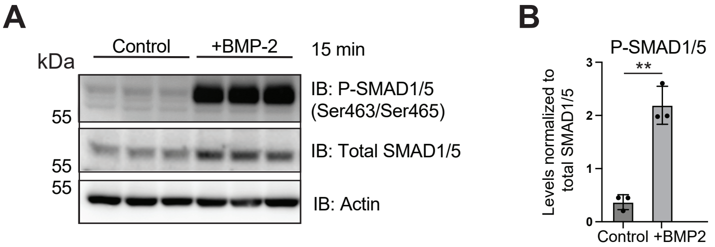


**Extended Data Fig. 12 | Cytokine treatment *in vitro*.**
**A.** Western blot analysis of phospho-SMAD1/5 (P-SMAD1/5), total SMAD 1/5 and actin in untreated (control) and BMP2 treated MVT1-GFP-luciferase cells. Treatment time amounted to 15 minutes. **B.** Bar graph showing **P**-SMAD1/5 levels normalized to total SMAD1/5, revealing activation of SMAD1/5 following BMP2 treatment. **, $P < 0.01$ by two-sided *t* test.
